# The *L. pneumophila* effector PieF modulates mRNA stability through association with eukaryotic CCR4-NOT

**DOI:** 10.1101/2022.06.06.494580

**Authors:** Harley O’Connor Mount, Francesco Zangari, Anne-Claude Gingras, Alexander W. Ensminger

## Abstract

The eukaryotic CCR4-NOT deadenylase complex is a highly conserved regulator of mRNA metabolism that influences the expression of the complete transcriptome, representing a prime target for a generalist bacterial pathogen. We show that a translocated bacterial effector protein, PieF (Lpg1972) of *L. pneumophila* Str. Philadelphia-1, interacts specifically with the CNOT7/8 nuclease module of CCR4-NOT, with a dissociation constant in the low nanomolar range. PieF inhibits the catalytic deadenylase subunit CNOT7 of the CCR4-NOT complex in a stoichiometric, dose-dependent manner *in vitro*. In transfected cells, PieF can silence reporter gene expression and reduce mRNA steady-state levels when artificially tethered. PieF demonstrates molecular similarities to another family of CNOT7-associated factors but demonstrates divergence concerning the interaction interface with CNOT7. In addition, we show that PieF overexpression changes the subcellular localization of CNOT7 and displaces the CNOT6/6L nucleases from CCR4-NOT. Finally, PieF expression phenocopies knockout of the CNOT7 ortholog in *S. cerevisiae*, resulting in 6-azauracil sensitivity. Collectively, this work suggests that *L. pneumophila* targets host mRNA stability and expression through a highly conserved host pathway not previously associated with *Legionella* pathogenesis.

## INTRODUCTION

*Legionella pneumophila* is a Gram-negative bacterial species found across a plethora of freshwater environments (1, 2). In nature, *L. pneumophila* replicates intracellularly within diverse protozoan hosts (3–5). Humans are typically exposed to *L. pneumophila* through inhalation of microbe-laden water droplets (6, 7) from contaminated water infrastructure resulting in Legionnaires’ disease (8–14). When *L. pneumophila* encounters human lung alveolar macrophages, a pathogenic program is initiated that is highly similar to that used within protozoan hosts (15–18). In both cases, the bacterium relies on the injection of pathogenic proteins, called ‘effectors’, through a type IV secretion system (19–22). Collectively, over 300 effectors prevent phagolysosome fusion with the *Legionella*-containing vacuole (LCV) and recruit host factors and membrane constituents to help nourish and accommodate the growing intracellular bacterial population (23). Individually, most effectors are non-essential for intracellular replication, due largely to genetic redundancy (24–26).

The *L. pneumophila* effector arsenal is encoded throughout its genome, comprising ∼10% of all open reading frames (27). Some effectors are encoded in regions of genomic plasticity that demonstrate variation between *L. pneumophila* strains (28–30). One such plasticity region was previously shown to encode several effectors that vary between *L. pneumophila* isolates and was subsequently named the **<u>p</u>**lasticity **i**sland of **<u>e</u>**ffectors (or *pie* locus) (30). The effectors within the *pie* locus are named *pieA-G* and are interspersed with several non-translocated bacterial genes as well (30). Since the discovery of this locus, PieA, PieE, and PieG have been characterized mechanistically (30–32). PieG (LegG1) is the best characterized and is prenylated at its C-terminus enabling localization to the LCV where it facilitates LCV motility through activation of host Ran GTPase (32–35). PieE was shown to associate with mammalian Rab GTPases (Rab5 and Rab7) and likely influences vesicular trafficking and GTPase recruitment to the LCV (31). The function of PieA is less obvious, but it localizes to the LCV as well (30).

Rewiring and silencing host gene expression are often critical mechanisms for the survival and replication of intracellular bacterial pathogens (36). Once inside a host cell, these invaders seek to dampen immune and/or stress response signalling by blocking transcription and translation of host defense factors (e.g., proinflammatory cytokines, pro-apoptotic caspases, positive and negative regulators of immune signaling pathways) (36–42). In addition to escaping targeting by the host, by limiting host gene expression, the pathogen can also free up micronutrients like amino acids that may be redirected for bacterial use (43–45). The mechanisms that pathogens use to modulate host gene expression are diverse and pathogen-dependent (46). *L. pneumophila* utilizes at least eight pathogenic factors to interfere with host translation through diverse mechanisms to alter the cell cycle and influence immune signaling cascades (39, 47). The effectors Lgt1-3 can mono-glucosylate the GTPase domain of eukaryotic elongation factor eEF1A, impairing function and translational elongation (48, 49). The effector SidI also binds eEF1A and eEF1β and impairs translation through uncharacterized mechanisms (50, 51). LegK4 phosphorylates host Hsp70 (T495) to impair ATPase, and chaperoning activity, driving aberrant polysome association (52, 53). Finally, *L. pneumophila* can silence host translation by blocking mTOR signaling through ubiquitination of upstream Rag GTPases (SidE) (45), or through uncharacterized mechanisms (SidL, RavX) (39, 41). This multifaceted and redundant approach to modulating host gene expression maximizes bacterial fitness within individual hosts, and across diverse host, types separated by millions of years of evolution (5, 26). The combined deletion of all effectors above did not alleviate translational repression during infection (45), suggesting that there are additional mechanisms employed to control eukaryotic gene expression yet to be described.

The eukaryotic CCR4-NOT complex is a large (>1 MDa) evolutionarily conserved molecular machine with critical roles in mRNA metabolism (54). The best-described role of CCR4-NOT is to shorten poly(A) tails of mature mRNA, as one of two major deadenylase complexes of eukaryotic cells (54). The other major deadenylase complex, PAN2/3, initially shortens mRNA poly(A) tails, priming them for the CCR4-NOT complex which further deadenylates the transcript, promoting either translation repression or decay pathways (for reviews see (54–57)). CCR4-NOT influences eukaryotic gene expression in concert with other deadenylases and CCR4-NOT-associated factors, allowing for temporal control of mRNA stability and turnover of gene expression programs (54). As a central regulator of eukaryotic gene expression, the CCR4-NOT complex has a critical role in ensuring proper cellular function (54). Given its vital cellular role, it is unsurprising that some subunits of CCR4-NOT are essential for mammalian development and cell viability (58).

The CCR4-NOT complex is scaffolded by the large multi-domain protein CNOT1 (59, 60). The MIF4G (Middle domain of eukaryotic initiation factor 4G) domain is found at the center of CNOT1 and is required for binding the DEDD (Asp-Glu-Asp-Asp) nuclease module (CNOT7 and CNOT8 in mammalian cells) (59, 60). CNOT7 binds the MIF4G domain in a mutually exclusive manner with CNOT8 and each subunit carries out similar, but distinct catalytic roles (61). In addition to the DEDD nuclease, CCR4-NOT contains a secondary endonuclease-exonuclease-phosphatase (EEP) nuclease module composed of the paralogs CNOT6 and CNOT6L (62, 63), which directly depend upon the DEDD module for incorporation into the CCR4-NOT complex (64, 65). Despite the catalytic similarity, each nuclease module regulates distinct mRNA subsets (66), with specific poly(A)-binding protein (PABP) occupancy preference (67). CCR4-NOT activity can be directed through the activity of adapter proteins including the antiproliferative BTG/TOB factors that associate with RNA-binding proteins (e.g., PABPC) to drive CCR4-NOT recruitment to mRNA (68–72).

Here we show that the *L. pneumophila* effector PieF (Lpg1972) binds to and modulates the activity of the host CCR4-NOT deadenylase complex. Specifically, PieF binds CNOT7/8, inhibits nuclease activity *in vitro*, and destabilizes reporter mRNA in transfected cells through apparent recruitment of CCR4-NOT. Taken together, this reveals CCR4-NOT modulation as another facet of highly conserved eukaryotic biology that is co-opted during *L. pneumophila* pathogenesis.

## MATERIAL AND METHODS

### Bacterial strains and culture conditions

Plasmids and *E. coli* strains used in this study are listed in Supplementary Table S1 and Table S2 respectively. All *L. pneumophila* strains used in the study (Table S3) were derived from Lp02 and Lp03 (73) and cultured as previously described (74). Briefly, *L. pneumophila* was grown at 37°C in N-(2-acetamido)-2-aminoethanesulfonic acid (ACES) buffered yeast extract supplemented with 100 μg/ml thymidine (AYET) and on charcoal AYET (CYET) agar plates. Liquid cultures were inoculated with patches grown for 2 days on CYET agar plates. Deletions were made as previously described (75), using primers listed in Supplementary Table S4.

### Yeast strains and culture conditions

Cultures were grown in either rich yeast peptone dextrose (YPD) medium (2% bacto peptone w/v, 1% yeast extract w/v, 2% glucose v/v), or synthetic defined medium (6.9 g/L of Formedium CYN0905) lacking specific amino acids where necessary. For solid media, agar was added to a final concentration of 2%. Cultures were grown overnight with shaking at 200 rpm at 30°C. *S. cerevisiae* strains used in this study are listed in Supplementary Table S5.

### Protein purification with 6X-His-Streptavidin binding peptide tag

Purifications were performed according to the QIAexpressionist protocol and as described (76). CNOT7 was amplified from pT7-EGFP-C1-HsNOT7 (Addgene #: 37325) (oHM478-479) and cloned into the T7 expression plasmid pMCSG68-SBP (Addgene #: 36943) resulting in an N-terminal 6xHis-SBP-TEV tag. PieF (aa1-125) and Lpg2149 (aa1-111) were cloned into pMCSG68-SBP. Briefly, *E. coli* BL21-Gold (DE3) chemically competent cells were transformed with each construct and grown in 1 L of LB broth with 100 μg/ml carbenicillin. Cultures were grown to an optical density (OD600nm) of 0.6 at 37°C, and expression was induced with 1 mM isopropylbeta-D- thiogalactopyranoside (IPTG) for 5 hours at 37°C. Cells were then harvested by centrifugation (12, 227 x *g* for 10 mins at 4°C). After resuspension in 40 ml lysis buffer (300 mM NaCl, 5% glycerol, 5 mM imidazole, 50 mM HEPES [4-(2-hydroxyethyl)-1-piperazineethanesulfonic acid] pH 7.5) cells were lysed by sonication on 30% amp (10 sec on, 10 sec off for 10 minutes) in the presence of 1 mM phenylmethylsulfonyl fluoride (PMSF). The lysate was then subjected to centrifugation (34, 957 x *g* for 20 mins at 4°C). The soluble fractions were then subjected to immobilized metal affinity chromatography purification with 2 ml nickel-nitriloacetic acid resin. Columns were washed with 300 ml wash buffer (300 mM NaCl, 30 mM imidazole, 15 mM HEPES pH 7.5, 5% glycerol), and eluted in 40 ml of elution buffer (300 mM NaCl, 300 mM imidazole, 15 mM HEPES pH 7.5). Eluates were dialyzed, quantified, and concentrated in dialysis buffer to >5 mg/ml (15 mM HEPES pH 7.5, 300 mM NaCl, 1mM dithiothreitol [DTT]) using Vivaspin 10 kDa cutoff columns (GE Healthcare 28932360). The 6X-His-tag was cleaved from CNOT7 for biolayer interferometry by incubating the protein with 6x- His-tagged TEV protease (60 μg/mg) overnight at 4°C. TEV protease and uncleaved protein were then removed by nickel resin purification, with the eluate kept for downstream assays.

### Identification of PieF host interacting proteins by AP-MS

To identify putative effector-host target interactions, we used the affinity purification coupled with mass spectrometry (AP-MS) approach as described (77) for two individual technical replicates. 6x-His-SBP PieF was incubated with magnetic streptavidin beads (Life Technologies) for 60 minutes in affinity purification buffer (50 mM HEPES pH 8, 150 mM NaCl, 1 mM EDTA, 0.2% NP-40). Human cell lysate from 5×10^7^ U937 cells was prepared through freeze-fracture lysis in affinity purification buffer with protease inhibitor cocktail (Roche P8340). The lysate was pre-cleared with streptavidin-conjugated beads to remove biotin-conjugated proteins. The lysate was further clarified through centrifugation (14, 000 x *g* for 15 min at 4°C). Streptavidin-bound PieF was then incubated with cell lysate for 3 hours at 4°C, purified through magnetic separation, and resuspended in 50 mM ammonium bicarbonate pH 8.0. Proteins were eluted by incubating with 2.5 mM biotin for 15 mins and digested overnight at 37°C with sequencing grade trypsin (Promega V511A). Trypsin digestion was terminated through the addition of trichloroacetic acid (0.5% final). Peptides were purified using C18 OMIX tips (Agilent AS57003) and eluted from the C18 tips using 0.1% formic acid in acetonitrile and dried by vacuum centrifugation. Before mass spectrometry the peptides were resuspended in 0.1% formic acid in water, centrifuged, and loaded in technical duplicate using an EASY-nLC II autosampler onto a 10 cm C18 column with a glass nano-spray ionization tip using EASY-nLC II liquid chromatography system (Thermo). For data collection, a Q-Exactive spectrometer was used in positive ion mode for a 120-min gradient on increasing acetonitrile concentration. Raw data files were converted using MSconvert and queried against the human and *L. pneumophila* proteome collection using GPM/X! tandem (http://www.thegpm.org/). Data were then analyzed using ProHits (78) by comparing the experiment to 20 unrelated AP-MS experiments with 11 unique baits as background controls. Filtering metrics included a GPM expect score of > -10, and < 8 unique peptides. The ribosomal, keratin, and artifact protein biofilters were also applied and excluded prey are listed in table S7. Data was sorted by unique peptides identified and normalized to protein size in kDa.

### Yeast two-hybrid assays

Yeast two-hybrid experiments were performed similarly to those described (79). Bait and prey proteins were fused N-terminally to the DNA-binding, and transcriptional activation domains of the Gal4p transcription factor respectively. This was achieved through Gateway cloning (80, 81) into pDEST-AD-ccdB or pDEST-DB-ccdB vectors, and then yeast transformation into RY1010 (MATa) or RY1030 (MATα) respectively (82). Yeast mating was performed as described (83) and diploid cultures were frozen in 25% glycerol for future use. To query for physical interactions between AD- and DB- fusions, diploids were grown overnight (SD -Leu -Trp) and diluted in 1 ml ddH2O to OD600nm=1. These dilutions were then serially diluted 5-fold to make 4 separate dilutions or used only at OD 0.04 as in Fig. 1C. 120 μL of each dilution as well as the undiluted culture were plated into 96-well plates. The diluted cultures were then stamped using the VP 407AH pin tool (V&P Scientific) onto either Y2H- selective (SD -Leu -Trp -His) or diploid selective media (SD -Leu -Trp) as in Fig. 3, 4A. Plates were incubated at 30°C and imaged 48 hours post-stamping. Disruptive yeast-two hybrid experiments were performed with a third vector pAG416-GPDp-ccdB which contains a Ura auxotrophic marker and constitutive GPD promoter (84). Y2H diploid strains were made chemically competent by subculturing to S-phase (OD600nm=2) and transformed with each respective vector. Assays were performed identically to above on diploid selective (SD -Leu -Trp -Ura) or Y2H-selective media (SD -Leu -Trp - Ura -His).

### Bacterial two-hybrid assays

B2H assays were performed similarly to those described in Karimova *et al.* (85). Each unique open-reading frame (ORF) was amplified with oligos designed using the NEBuilder software (V2.4.0) (Table S4) for use with the pKT25 and pUT18C B2H vectors. Amplicons were incubated with XbaI linearized vector according to the manufacturer’s recommendations for the NEBuilder HiFi DNA assembly cloning mix. Co-transformants were selected on LB media supplemented with antibiotics (50 μg/ml kanamycin [KAN], 100 μg/ml carbenicillin [CARB]). Cultures of individual co-transformant clones were grown to saturation overnight at 37°C in LB (50 μg/ml KAN, 100 μg/ml CARB). and then diluted to OD600nm=0.2 in ddH2O. The dilutions were then pinned as described above onto supplemented LB (50 μg/ml KAN, 100 μg/ml CARB, 0.5 mM IPTG, 40 μg/mL 5-Bromo-4-chloro-3- indolyl- β-D-galactopyranoside (X-gal)), as well as supplemented M63 media with maltose (25 μg/ml KAN, 50 μg/ml CARB, 0.5 mM IPTG). Plates were incubated for 3-5 days at 30°C and imaged using an S&P robotic imager.

### Biolayer interferometry binding assays

Assays were performed on the Octet RED96 platform (PALL-ForteBio). Sensor-bound samples were diluted to 5 μg/ml in kinetics buffer for each assay (1× PBS, 0.1 mg/ml bovine serum albumin, 0.002% Tween 20) for each assay. Streptavidin biosensors were equilibrated in kinetics buffer and then moved to purified tagged PieF solution at 5 μg/ml or kinetics buffer alone. Following a 200-second loading period, the sensors were again washed for 450 seconds in kinetics buffer and incubated with a range of concentrations of untagged catalytically active CNOT7 (0, 27, 54, 108, 270, 540, 1080 nM). Following an association period of 300 seconds, the sensors were again moved to kinetics buffer for 450 seconds to allow dissociation of CNOT7. Data analysis was performed using the GraphPad Prism software (Prism 9). Data was fitted through non-linear regression using a one-site-specific binding model. Each measurement was performed a minimum of three times in independent experiments.

### Deadenylation assay

Purified protein was diluted in 25 μL deadenylase buffer (50 mM Tris-HCl pH7.5, 10% glycerol, 1 mM DTT, 2 mM MgCl2, 150 mM NaCl). PieF, CNOT7, and Lpg2149 were pre-incubated at room temperature in 50 μL reactions. Lpg1972 nuclease inhibitory activity was tested at a range of decreasing concentrations including (1, 0.5, and 0.25 μM as determined by nanodrop). CNOT7 was included at 1 μM final concentration. Following pre-incubation, 2.5 μL of PAGE-purified 10 μM 5′-6 carboxyfluorescein labelled RNA substrate with a poly(A) tail of 20 nucleotides [5′-FAM- UCUAAAUAAAAAAAAAAAAAAAAAAAA-3′] (IDT) was added to each reaction and incubated at 37°C for 30 min. Reactions were stopped with the addition of RNA loading buffer II (Thermo Fisher AM8546G) and boiling at 95°C for 5 min. 5 μL of each reaction was loaded onto an 8M UREA 25% polyacrylamide gel and electrophoresed at 180V. The gel was then fixed in 1X TBE with 10 % acetic acid and 10% methanol for 5 min at RT. The fixed gel was imaged on the Typhoon (FLA 9500) at 800 V using a 488 nm excitation wavelength. The gel image is representative of at least three independent experiments.

### λN Tethering assay

HEK293T cells were seeded in 24-well plates (Eppendorf 0030722116) at 2×10^5^ cells per well. Cells were transfected using the Lipofectamine 2000 manufacturers protocol (800 ng endotoxin-free DNA at a 1:1:1 ratio and 2 μL Lipofectamine). After 16-20 hours post-transfection cells were lysed in passive lysis buffer for 1 hour (Promega E1910) and processed using the Promega Dual-Luciferase Reporter Assay System with 2.5 μL of lysate (Promega E1910). Luminescence was measured using a TECAN infinite 200 pro Mplex machine with a 10, 000 ms integration time. Luciferase activity results represent three independent biological replicates. For steady-state mRNA measurements, RNA was extracted from cells prepared as above according to the manufacturer’s protocol (PureLink RNA mini kit 12183018A). On-column DNase digestion was performed according to the manufacturer’s protocol (Ambion PureLink DNase set 12185010). RNA was quantified by nanodrop and 1 μg was used for cDNA synthesis using the SuperScript Vilo cDNA synthesis kit (ThermoScientific 11754050). qRT-PCR was performed with 1 μL template, to determine reporter steady-state levels for *Renilla* and Firefly luciferase on a Biorad CFX96 machine. *Renilla* Ct values were normalized relative to firefly luciferase, and dCT values were used for the Student’s t-test of significance. dCT values were normalized relative to LacZ negative control samples and plotted. Steady-state level plots represent measurements of two technical replicates and are representative of at least three biological replicates.

### Mammalian stable cell line construction for mass spectrometry of CCR4-NOT composition

To generate stable cell lines, the Flp-In™ T-REx™ 293 (ThermoFisher R78007) cells were transfected according to the Lipofectamine 2000 protocol (ThermoFisher 11668027). On day 0 cells were seeded in a 6-well plate at 40% confluency and grown overnight at 37°C and 5% CO2. On day 1 cells were transfected with 2.5 μg of pOG44, and 200 ng of each 3′ 3X-FLAG-tagged CNOT7, or empty FLAG vector in 250 μL Opti-MEM (ThermoFisher 31985062) with 10 μL of Lipofectamine 2000. On day 2 cells were re-plated to 10 cm dishes in Dulbecco’s Modified Eagle Medium (DMEM) with 10% fetal bovine serum (FBS). On day 3 hygromycin B was added to a final concentration of 200 μg/mL to select for stable integrants. DMEM media was changed every three days. Mammalian cell lines used in this study are listed in Table S6.

### Transfection of HA-mCherry vectors into stable cell lines for AP-MS

Stable cell lines were generated using the Flp-In™ T-REx™ 293 system (ThermoFisher R78007) by seeding a 6-well plate with 4.8×10^5^ cells/ well. The following day cells were transfected with 2.5 μg pOG44, 200 ng pcDNA 3′ FLAG-CNOT7 in 250 μL Opti-MEM (ThermoFisher 31985062) with 10 μL of Lipofectamine 2000. On day 2, cells were re-plated to 10 cm dishes. On the third day hygromycin, B was added to a final concentration of 200 μg/ml to select for stable integrants. DMEM media was changed every three days until the cells were confluent. For CNOT7 induction before FLAG AP-MS, cells were seeded and grown to approximately 60-70% confluency in DMEM medium supplemented with 10% fetal bovine serum (Wisent 098150) in 15 cm dishes. Cells were then induced with 1 μg/ml of tetracycline (BioBasic TB0504) and transfected with 145 μL Lipofectamine 2000 (ThermoFisher 11668027) containing 58 μg of HA-mCherry tagged effector DNA as described above. After 24 hours cells were visually inspected for fluorescence by microscopy (Leica DMi8 inverted fluorescent microscope) to confirm the expression of each effector before harvest. Cells were harvested by washing once with 10 ml of PBS and pelleting at 100 x *g* for 5 minutes. The cells pellets were frozen in liquid nitrogen, weighed, and stored at -80°C until further processing.

### Immunoprecipitation of FLAG-tagged CNOT7 and preparation for mass spectrometry

Cell pellets from three independent 15 cm dishes were prepared as biological replicates. Pellets were lysed at 4°C in lysis buffer (50 mM HEPES-NaOH (pH 8.0), 100 mM NaCl, 2 mM EDTA, and 10% glycerol with freshly added 0.1% NP-40, 1 mM DTT, 1 mM PMSF, and protease inhibitor cocktail (Sigma-Aldrich P8340, 1:500). Lysis buffer was added in a 1:4 pellet weight to volume ratio (0.4 ml of lysis buffer per 0.1g pellet). Freeze fracturing was performed by freezing tubes in a dry ice ethanol bath and then thawing slowly in a 37°C water bath. After thawing samples were placed back on ice and sonicated with three bursts at an amplitude of 25% for 5 s with 3s of rest in between using a Q500 Sonicator with a 1/8” microtip (Qsonica, Newton, Connecticut, Cat#4422). Samples were then digested with 250U benzonase to reduce nucleic acid-mediated interactions (Sigma-Aldrich, E8263, 250U) for 30 min at 4°C with end-over-end rotation. Lysates were cleared at 20, 000 x *g* for 20 min at 4°C. Clarified lysates were normalized by volume and incubated with 60 μL of a 33% magnetic anti-FLAG M2 bead slurry (20 μL beads) (Sigma-Aldrich, M8823) equilibrated in lysis buffer.

Immunoprecipitation proceeded for 3 hours at 4°C with end-over-end rotation. Following incubation, the magnetic beads were pelleted at 1000 x *g* for 30 seconds at 4°C, magnetized and lysate was removed. The beads were then washed with 1 ml of lysis buffer at 4°C and transferred to new tubes. Beads were washed again with 1 ml lysis buffer and then washed once more with 1 ml ammonium bicarbonate buffer (50 mM, pH 8.0). After washing, the beads were resuspended in 10 μL ammonium bicarbonate (50 mM, pH8.0) with 1000 ng trypsin (Sigma-Aldrich, T6567) and incubated at 37°C overnight, with rotation, Beads were magnetized, and the supernatant was collected. The second round of digestion on beads was performed with 500 ng of trypsin in 5 μL of ammonium bicarbonate buffer (50 mM, pH 8.0) and rotated for 4 h at 37°C. Digestion was stopped by the addition of formic acid to a final concentration of 2.5%.

### Mass spectrometry acquisition using TripleTOF mass spectrometers

AP-MS samples were subjected to mass spectrometry in three biological replicates. A quarter of each sample was loaded at 800 nL/minute onto a 15 cm 100 μm ID emitter tip packed in-house with 3.5 μm Reprosil C18 (Dr. Maisch GmbH, Germany). The peptides were eluted from the column at 400 nL/min over a 90-minute gradient generated by a 425 NanoLC (Eksigent, Redwood, CA) and analyzed on a TripleTOF™ 6600 instrument (AB SCIEX, Concord, Ontario, Canada). The gradient started at 2% acetonitrile with 0.1% formic acid and increased to 35% acetonitrile over 90 minutes followed by 15 minutes at 80% acetonitrile, and then 15 min at 2% acetonitrile for a total of 120 minutes. After each sample, the column was flushed twice for one hour at 1500nL/min using an alternating sawtooth gradient from 35% acetonitrile to 80% acetonitrile. Each concentration gradient was maintained for five minutes. Column and instrument performance was verified after each sample by also analyzing a 30 fmol bovine serum albumin (BSA) tryptic peptide digest with a 60 fmol α-casein tryptic digest in a short 30-minute gradient. Mass calibration for the mass spectrometer was performed on the BSA reference ions between samples. Acquisition of ions was performed in data-dependent mode and consisted of one 250 ms MS1 TOF survey scan from 400-1250 Da followed by twenty 100 ms MS2 candidate ion scans from 100–2000 Da in high sensitivity mode. Ions with a charge of 2+ to 4+ exceeding the threshold of 200 counts per second were selected for fragmentation. Former precursors were excluded for 10s after one occurrence,

### MS peptide analysis and SAINT filtering

Data from the mass spectrometer was analyzed using the ProHits laboratory information management system (LIMS) platform (78). Wiff files were converted to MGF format using WIFF2MGF. MGF files were then converted to mzML format with ProteoWizard (3.0.4468) and AB SCIEX MS Data Converter (V1.3 beta), MzML files were searched using Mascot (v2.3.02) and Comet (2014.02 rev.2) (86). Search results were concatenated and analyzed with the Trans-Proteomic Pipeline (TPP) via the iProphet pipeline. The spectra were searched against a total of 72, 518 proteins consisting of the NCBI human RefSeq database (version 57; 27 March 2020; forward and reverse sequences), *Legionella pneumophila subsp. pneumophila* (strain Philadelphia 1 / ATCC 33152 / DSM 7513) Lpg1972, Lpg2137 and Lpg2149 along with common contaminants from MaxQuant (87) and the Global Proteome Machine (http://www.thegpm.org/crap/index.html), as well as sequences from common fusion proteins and epitope tags. Database parameters were adjusted to search for tryptic cleavages, allowing up to two missed cleavage sites per peptide, MS1 mass tolerance of 40 ppm with charges of 2+ to 4+ and an MS2 mass tolerance of ± 0.15 amu. Asparagine/glutamine deamidation and methionine oxidation were selected as variable modifications. A minimum iProphet probability of 0.95 was required for protein identification. Proteins detected with a minimal number of two unique peptides were used for protein interaction scoring. Significance Analysis of INTeractome (SAINTexpress version 3.6.1) was applied to calculate the probability of potential interactions from the background (88). Three biological replicates of a cell line expressing only the FLAG tag were subjected to the same process as FLAG-CNOT7 cell lines to function as a negative control. Each biological replicate was analyzed independently against the control before averaging biological replicate results and calculating Bayesian False Discovery Rates (BFDR). High-confidence interactions were those with BFDR ≤ 1%. Data has been deposited as a complete submission to the MassIVE repository (https://massive.ucsd.edu/ProteoSAFe/static/massive.jsp) and assigned the accession number MSV000089325. The ProteomeXchange accession is PXD033482.

### *S. cerevisiae* kinetic growth curve assays

Yeast cultures were grown to saturation in 3 ml of synthetic defined media (SD-Ura). The culture optical density (OD600) was then measured, and 5 ml of inoculum was prepared at an OD600 of 0.1. Serial two-fold dilutions of analyte (6-azauracil [Sigma-Aldrich A1757-5G]) were made along the x-axis of a 96 well plate in 100 μL volumes. Wells were inoculated with 100 μL inoculum preparation and sealed with a breathable membrane (Sigma Z380059-1PAK). The plates were then incubated at 30°C using an S&P growth curve robot, with 15-minute read intervals at OD620. Results were plotted for individual analyte concentrations using the R package ggplot2 (V3.3.5). Benjamini-Hochberg corrected Student’s t-tests were performed to assess statistical significance. Results are representative of at least two biological replicates.

### Fluorescence microscopy

Imaging was performed using a Leica DMi8 inverted fluorescent microscope. HEK293T cells were transfected using Lipofectamine 2000 according to the manufacturer’s protocol (800 ng DNA, 2 μL Lipofectamine 2000) and then seeded at 1×10^5^ cells on poly-L-lysine coated glass coverslips in 24- well plates immediately following transfection. After 16-20 hours post-transfection cells were washed gently three times with PBS. Cells were then fixed in 4% paraformaldehyde (Electron Microscopy Sciences 15710) for 20 min at RT and washed three more times with PBS. The cells were then permeabilized in PBS + 0.5% Triton X-100 for 5 mins at RT. The cells were again washed three times and then the coverslips were mounted onto microscope slides with Prolong GOLD Anti-fade with DAPI (Life Technologies P36935). The mounted slides were allowed to cure for 24 hours in the dark at RT. Nuclear exclusion statistics were calculated as a Student’s t-test between each sample.

## RESULTS

### The Dot/Icm substrate PieF directly binds CNOT7 of the CCR4-NOT complex with low nanomolar affinity

The *pie* locus encodes several effectors, named PieA-G (30), as well as several non-translocated bacterial proteins. PieF (Lpg1972) is the smallest effector in the Pie locus (Fig. S1A) and remains largely uncharacterized. Like most *L. pneumophila* effectors, PieF is dispensable for growth in U937 cells (Fig. S1B) and four protozoan hosts (89) and is only characterized by homology to coiled-coils domains (30). To investigate the function of PieF, we took an affinity purification coupled to mass spectrometry (AP-MS) approach (77) to identify potential host targets. Purified 6X-His-Streptavidin-Binding peptide (SBP) tagged PieF was immobilized on streptavidin magnetic beads and incubated with mammalian U937 cell lysate. Non-specific proteins were washed away before on-bead tryptic digest and MS identification. Co-precipitating peptides were filtered against 11 unrelated bait proteins (encompassing 20 unique effector control AP-MS experiments), as well as the keratin, ribosome, and artifact ProHits biofilters to eliminate common precipitation partners (Table S7). In total 12 prey proteins exceeded our detection threshold (an average of > 8 unique peptides supporting the prey identity). Notably, five components of the eukaryotic CCR4-NOT complex co-precipitated with PieF above background (CNOT1, CNOT2, CNOT3, CNOT7, CNOT9) (Fig. 1A and B). The CCR4- NOT-associated factor TNKS1BP1 also co-precipitated with PieF (61). Additionally, we observed precipitation of PAN3, a non-catalytic component in the general eukaryotic deadenylase machinery (90). The early cyclin-dependent kinases CDK4 and CDK6, known interactors of CCR4-NOT (91), were also detected, which could suggest a role for PieF in modulating the eukaryotic cell cycle. When peptide counts were normalized by protein size the top identified interactor was the DEDD deadenylase, CNOT7, from the CCR4-NOT complex (Figs. 1A and B).

**Figure 1.**
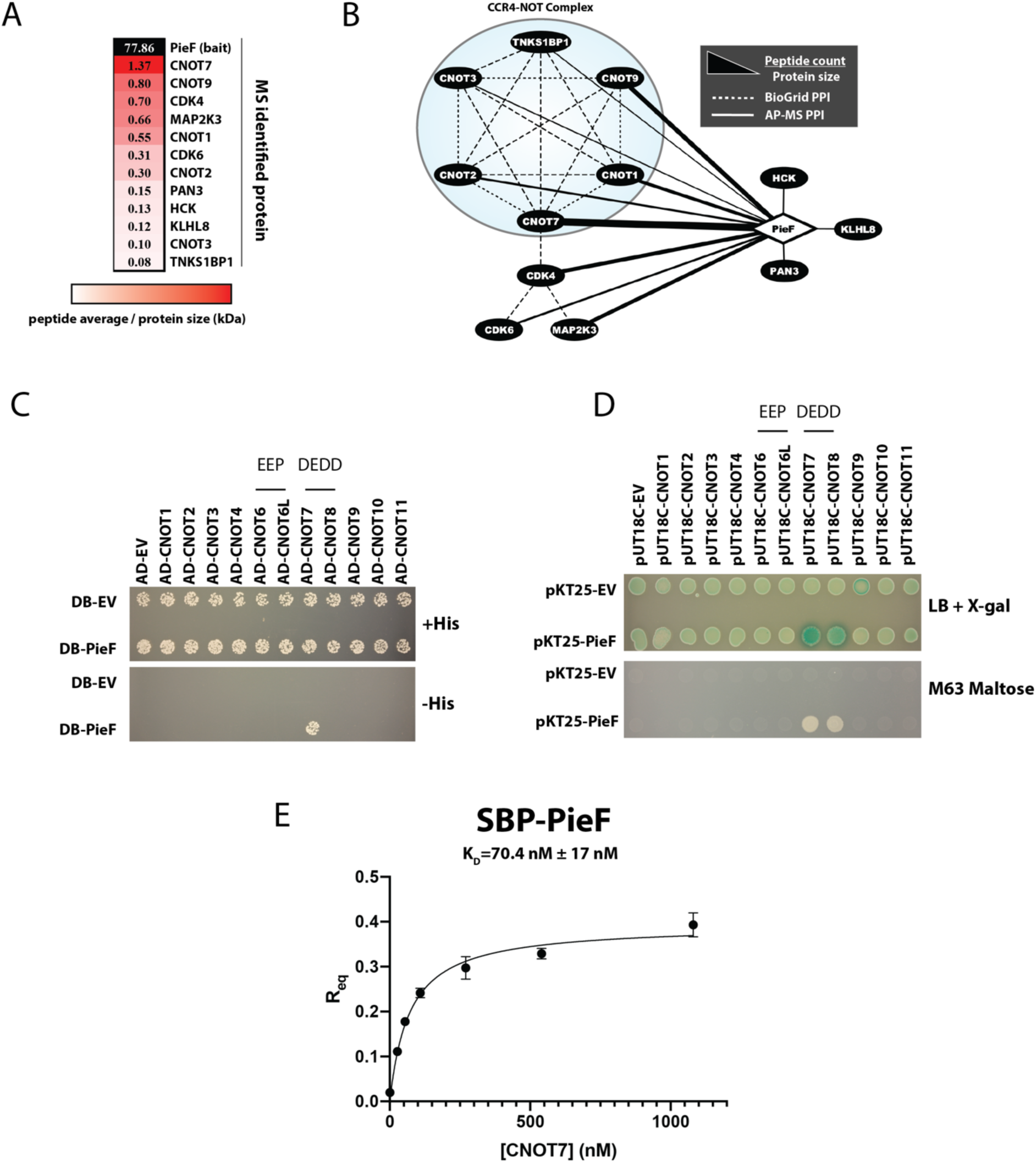
Identifying PieF host targets by AP-MS. To identify host proteins interacting with PieF, purified SBP-PieF was immobilized on streptavidin beads and incubated with U937 lysate. Proteins co-purifying with PieF were identified using mass spectrometry. **A)** A heatmap showing the normalized spectral counts (peptide count/protein size (kDa) of the bait PieF and co-purifying proteins. Prey with fewer than 8 average unique peptides supporting the interaction were filtered. **B)** AP-MS network representation of prey proteins that co-precipitate with PieF bait. Network edges represent peptide spectral counts normalized to protein size [spectral counts/size (kDa)]. BioGrid database (https://thebiogrid.org/) interactions are illustrated with dashed edges. CCR4-NOT complex enrichment encircled. **C)** Y2H assay demonstrating a likely direct physical interaction between PieF and the DEDD nuclease CNOT7. Transcription factor complementation stimulates *HIS3* expression allowing survival on media lacking histidine (-His). **D)** B2H assay demonstrating a likely direct physical interaction between pieF and both DEDD nuclease subunits CNOT7 and CNOT8. Adenylate cyclase complementation stimulates expression of the maltose operon, as well as LacZ induction. This complementation can be observed by growth on M63 with maltose as the sole carbon source, or through cleavage of 5-Bromo-4-chloro-3-indolyl-β-D-galactopyranoside (X-gal) by LacZ. **E)** Biolayer interferometry saturation curves acquired with a Forte-Bio instrument. Binding affinity measurements for CNOT7 against sensor bound His-SBP-PieF were fit using a single site-specific binding model in Prism (Prism 9). Results are representative of three independent experiments. Error bars represent the SEM of three independent experiments.

The orthogonal yeast two-hybrid (Y2H) assay (92) was used to confirm the physical interaction partner(s) identified in our AP-MS dataset. We tested each component of the human CCR4-NOT complex for a physical interaction with PieF given the enrichment observed for this complex during AP-MS. Only the DEDD nuclease subunit CNOT7 had a Y2H interaction with PieF (Figure 1C), consistent with our robust peptide enrichments for CNOT7 (Figs. 1A and B). The CCR4- NOT complex is widely conserved across eukaryotes, and this complex was originally described in *S. cerevisiae* (93). To rule out the possibility that an *S. cerevisiae* ortholog was bridging an indirect interaction between PieF and CNOT7 during Y2H, we next sought to confirm the interaction using the orthogonal adenylate cyclase-based bacterial two-hybrid assay (B2H) (85). The B2H approach is performed within *Escherichia coli,* which lacks endogenous CCR4-NOT components. We observed a detectable B2H signal, denoted by growth on M63 maltose and X-gal cleavage, between PieF and both DEDD nuclease subunits (CNOT7/8) (Fig. 1D). Given the high degree of primary sequence similarity between these paralogs (76% identical) (61), it is unsurprising that PieF is capable of interacting with both DEDD nuclease subunits if it binds either nuclease individually. However, as with all physical interaction detection methods, sensitivity limitations, may explain why an interaction with CNOT8 was not detected by AP-MS or Y2H (94).

Our data implicate a direct physical interaction between PieF and CNOT7 using three orthogonal detection methods. To measure the binding affinity between PieF and CNOT7, biolayer interferometry (BLI) kinetic measurements were gathered using a ForteBio Octet instrument with purified proteins. The dissociation constant of untagged CNOT7 from PieF was calculated at 70.37 nM (± 17 nM) (Fig. 1E). To rule out non-specific binding, we also performed a negative control experiment in which untagged CNOT7 was incubated with a bare streptavidin biosensor at the highest tested concentration (1080 nM). CNOT7 failed to bind the streptavidin biosensor in the absence of PieF (Table 1). Taken together, these results demonstrate that PieF binds CNOT7 (and CNOT8) protein with a biologically relevant affinity (95). The BTG/TOB family of proteins is known to interact with CNOT7/8 of CCR4-NOT (96), for context, the dissociation constant between PieF and CNOT7 was ∼5-9 times lower than reported between CNOT7 and TOB by isothermal titration calorimetry (97, 98).

**Table 1.** BLI raw data. Measurements from one BLI experiment illustrate that CNOT7 requires sensor bound PieF for a detectable BLI signal.

| **Sensor Bound**<br>**PieF** | **CNOT7 Conc.**<br>**(nM)** | **Response** |
| --- | --- | --- |
| Y | 1080 | 0.4323 |
| Y | 540 | 0.3416 |
| Y | 270 | 0.3181 |
| Y | 10 | 0.262 |
| Y | 54 | 0.1781 |
| Y | 27 | 0.1098 |
| Y | 0 | 0.0229 |
| N | 1080 | 0.0263 |

### The modulation of CNOT7 by PieF mimics eukaryotic BTG/TOB proteins

As CCR4-NOT regulates diverse aspects of eukaryotic biology including apoptosis, immune-signalling, and cellular proliferation, identifying a specific output of PieF-mediated modulation represents a significant challenge. After demonstrating a direct physical interaction between PieF and CNOT7, we next investigated whether PieF modulates CCR4-NOT activity in a variety of established assays. First, an *in vitro* deadenylase assay was performed with purified proteins. The best-characterized molecular activity of CNOT7/8 is to deadenylate mRNA transcripts as a 3′-5′ exonuclease (99, 100). Several studies have used purified DEDD, and EEP subunits of CCR4-NOT from bacterial expression systems to assay these proteins for *in vitro* nuclease activity against synthetic RNA substrates (97, 101–104). To assay whether the nuclease activity of CNOT7 is influenced by PieF, we repeated this *in vitro* approach in the presence and absence of purified PieF. Comparable to previous studies, purified CNOT7 showed robust deadenylase activity against a 5′ fluorescently-labelled (6-FAM) RNA substrate (97, 101–103), resulting in a range of product sizes (Fig. 2A). On their own, purified PieF and our negative effector control, Lpg2149, showed no deadenylase activity against this substrate. Lpg2149 was chosen as an effector control because it does not target host proteins during infection (105) but its molecular weight closely approximates that of PieF. When PieF was preincubated with CNOT7 at equimolar ratios nuclease activity was completely blocked (Fig. 2A). The inhibition of CNOT7 was specific to PieF and appears to be stoichiometric, diminishing at molar ratios below 1:1 (Fig. 2A). One explanation is that PieF may sterically occlude the active site of CNOT7 *in vitro*, although this is not the case for BTG/TOB proteins (96, 104, 106). Alternatively, PieF binding may promote conformational changes in CNOT7, triggering the release of the RNA substrate for another deadenylase (CNOT6/6L), or process (De-capping, etc.) (107).

**Figure 2.**
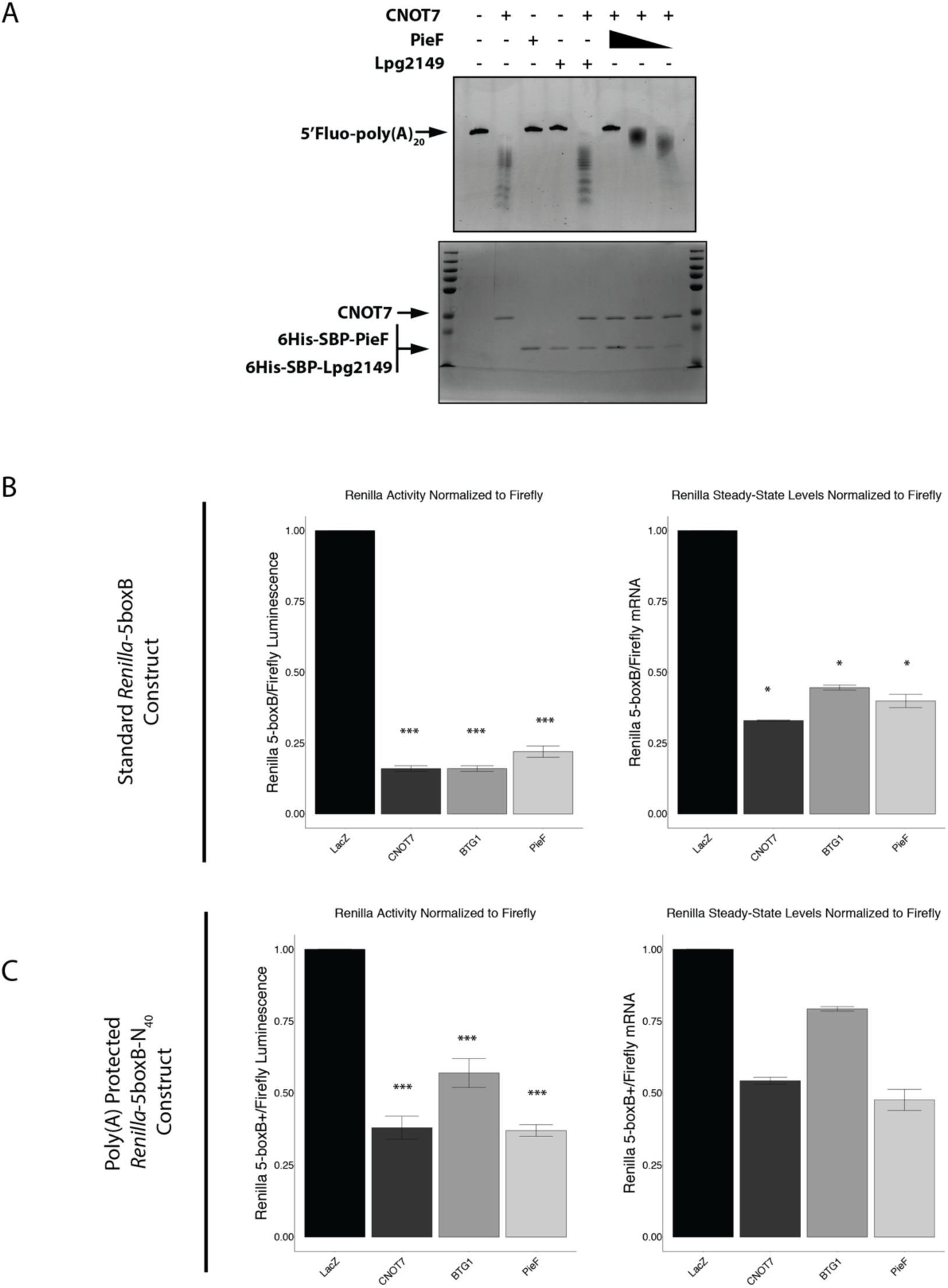
PieF phenocopies BTG molecular function. **A)** PieF can catalytically impair the degradation of a synthetic RNA substrate at equimolar ratios with CNOT7. The RNA substrate is labelled at the 5’ end with 6-carboxyfluorescein and contains a 20-nucleotide poly(A) tail. RNA products are resolved on 25% 8M UREA-PAGE (top gel). Proteins are resolved separately for loading controls on 10% SDS-PAGE stained with Coomassie Blue (bottom gel). PieF concentrations are 1 μM, 0.5 μM, 0.25 μM. All other proteins were assayed at 1 μM. **B)** Using a *Renilla* luciferase (RL) reporter construct with five BoxB hairpins within the 3’ UTR (RL-5BoxB), mRNA levels and activity are measured relative to a firefly (FL) control. Schematics are shown above data. **C)** Using a modified luciferase reporter construct with 5x BoxB hairpins and a protected poly(A) tail, mRNA levels and activity are measured relative to an FL control to assess the requirement of deadenylation on silencing. For B and C, the RL activity is normalized to FL activity for each sample, and then the ratio is plotted relative to lacZ. Activity data represents an average and standard deviation of three biological replicates. Results are representative of at least three biological replicates. Student’s t-test was calculated on the ratio of RL/FL luminescence to determine significance (* = p < 0.05, *** = p < 0.01). qRT-PCR results illustrate one representative biological replicate, average and standard deviation of two technical replicates plotted. Student’s t-test was calculated on RL dCT values normalized to FL (* = p < 0.05).

While PieF inhibits CNOT7 deadenylase activity *in vitro,* some endogenous modulators of the CCR4-NOT complex with inhibitory roles *in vitro* end up being more nuanced when examined in live cells. An example is BTG2 which inhibits CNOT7 deadenylase activity in a dose-dependent manner *in vitro* (107, 108), but in the cellular context directs the complex to degrade mRNA transcripts required for cellular proliferation (68-70, 91, 96, 109-113). Like PieF, BTG1/2 are small proteins with no catalytic core, or defined RNA-binding domains (107). As *L. pneumophila* is known to possess effector proteins that mimic the functions of eukaryotic proteins (28, 29, 114, 115), we sought to further compare the CCR4-NOT modulatory activities of PieF and BTG/TOB proteins.

An established methodology for studying CCR4-NOT activity within the cellular context is the artificial tethering of proteins to mRNA reporter transcripts (116). When tethered, components of the CCR4-NOT complex (117, 118), and BTG/TOB proteins (113, 118) destabilize reporter mRNA and reduce reporter gene expression in live cells. BTG/TOB-mediated destabilization is due to the recruitment of endogenous CCR4-NOT complex (68, 71, 72, 113). The *in vitro* (107, 108) and *in vivo* (68) discrepancies of BTG/TOB activity prompted us to assess the impact of PieF on RNA stability and expression. To this end, we leveraged the bacteriophage λN-BoxB tethering assay in HEK293T cells (119–121). Using this approach, we measured the impact of tethering CNOT7, PieF, or BTG1 to two different reporters: 1) a standard *Renilla* reporter with five BoxB hairpins within its 3′ UTR or 2) a modified *Renilla* reporter with five BoxB hairpins within its 3′ UTR as well as a self-cleaving hammerhead ribozyme (HHR) which generates a protected poly(A) tail that is not accessible to 3′-5′ exonucleases (121). The modified reporter allows for deadenylation-independent mechanisms of silencing to be assayed.

Under each condition the results with PieF or BTG1 were comparable: both proteins reduced reporter expression, and mRNA steady-state levels of the standard reporter, as does CNOT7 alone (Fig. 2B). This effect was not observed for the negative control LacZ. PieF and BTG1 could also reduce expression of the modified, poly(A) protected *Renilla* reporter, without significantly impacting steady-state mRNA levels (Fig. 2C). This effect was again observed with CNOT7 and not for LacZ. Using the modified reporter, BTG1 appeared to have a less inhibitory effect than CNOT7 or PieF alone. The cause of this effect is unclear but may indicate that PieF can more effectively recruit the CCR4-NOT complex when artificially tethered, consistent with comparisons between our affinity measurements and those reported for TOB1 (97, 98). This interpretation should be considered carefully, however, as it relies on comparisons of TOB-CNOT7 affinity, rather than BTG1-CNOT7 (97, 98). Together, these results support that PieF can recruit the CCR4-NOT complex to mRNA substrates, like the BTG/TOB protein class. CCR4-NOT recruitment by either BTG/TOB or PieF subsequently stimulates mRNA silencing and/or degradation. Furthermore, given that PieF can inhibit the expression of both the standard and modified reporters it suggests that silencing can occur through deadenylation-dependent and -independent mechanisms. Previously subunits of the CCR4- NOT complex, including catalytically active and inactive CNOT7, have illustrated translational silencing independent of deadenylation activity (122–124). The GW182 protein family (TNRC6A, B, C) of the miRNA silencing pathway also translationally represses targets through CCR4-NOT-mediated PABP dissociation that is not dependent on deadenylase activity (121).

### The binding of PieF to CNOT7 is unaffected by mutations that ablate the interaction with BTG1

Phenotypic comparisons between PieF and BTG/TOB proteins led us to explore whether these proteins interact with CNOT7 analogously. Through examination of co-crystal structures, CNOT7 residues required for BTG/TOB interaction (E247, Y260) have been identified and experimentally validated (125). When CNOT7 was mutated at both sites (E247A, Y260A), the Y2H interaction with BTG1 was abrogated, however, these mutations failed to disrupt the CNOT7-PieF interaction (Fig. 3). This result suggests that PieF associates through a different or expanded binding interface that does not require contacts with CNOT7 E247, or Y260.

**Figure 3.**
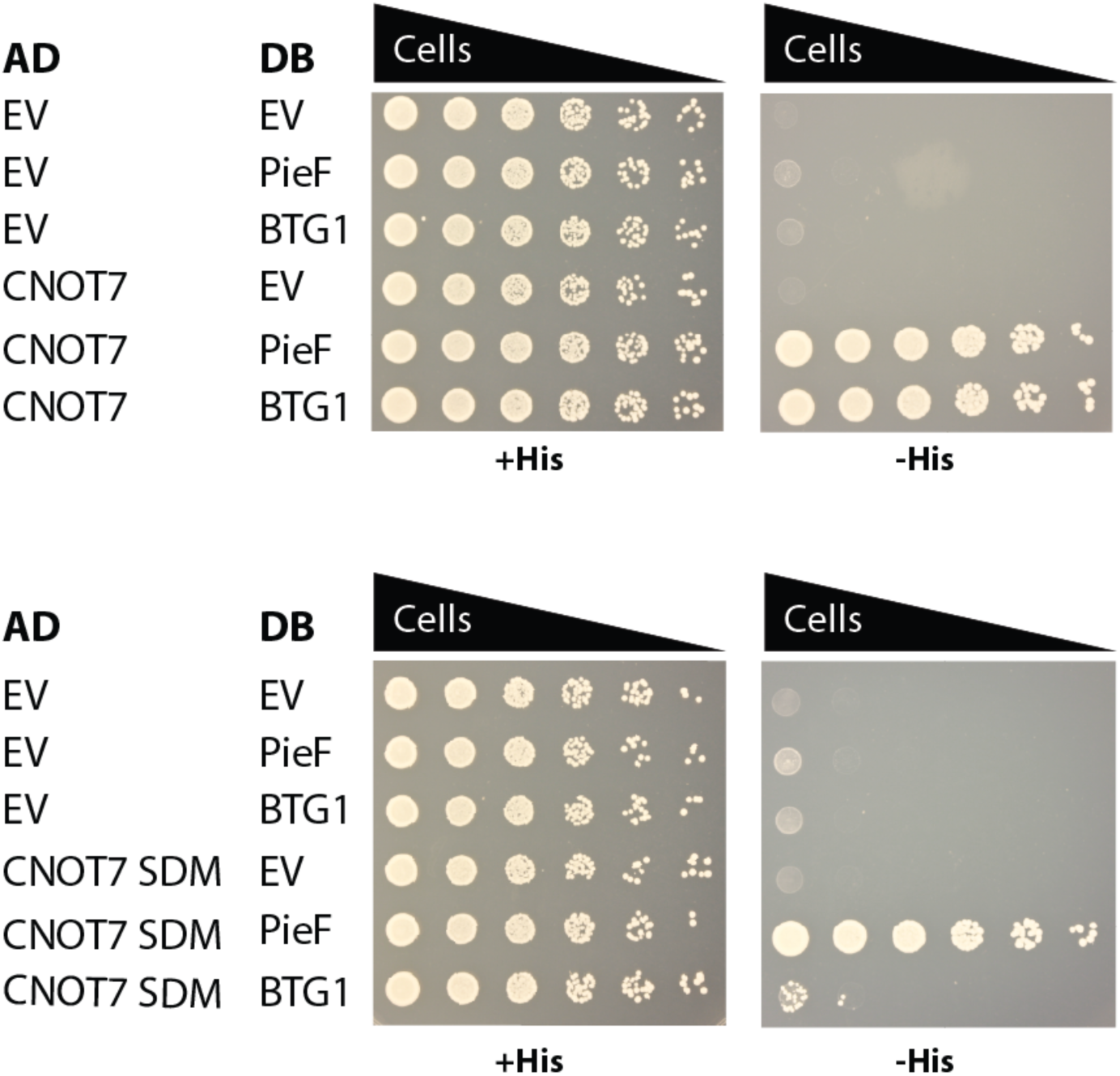
The binding of PieF to CNOT7 is unaffected by mutations that ablate the interaction with BTG1. Y2H spot-dilution was performed with undiluted culture (leftmost column #1) as well as serial 5-fold dilutions of culture from OD600=1.0 to 0.0016 (columns 2-6). A CNOT7 mutant (E247A, Y260A) was previously shown unable to interact with BTG2 (125). These mutations could also ablate the Y2H signal with BTG1, but not with PieF.

### PieF modulates CCR4-NOT subunit composition

Next, we examined the effect of PieF on CCR4-NOT complex subunit composition upon binding to CNOT7. To test this, we used a Y2H assay with a third untagged protein that was constitutively expressed from a low-copy vector to assess how the physical interaction between CNOT7 and CNOT6L may change in the presence of PieF or control Lpg2149. Expression of PieF, but not Lpg2149, was sufficient to diminish the Y2H signal between CNOT7 and CNOT6L (Fig. 4A). One proposed competitive binding model is illustrated (Fig. 4B).

**Figure 4.**
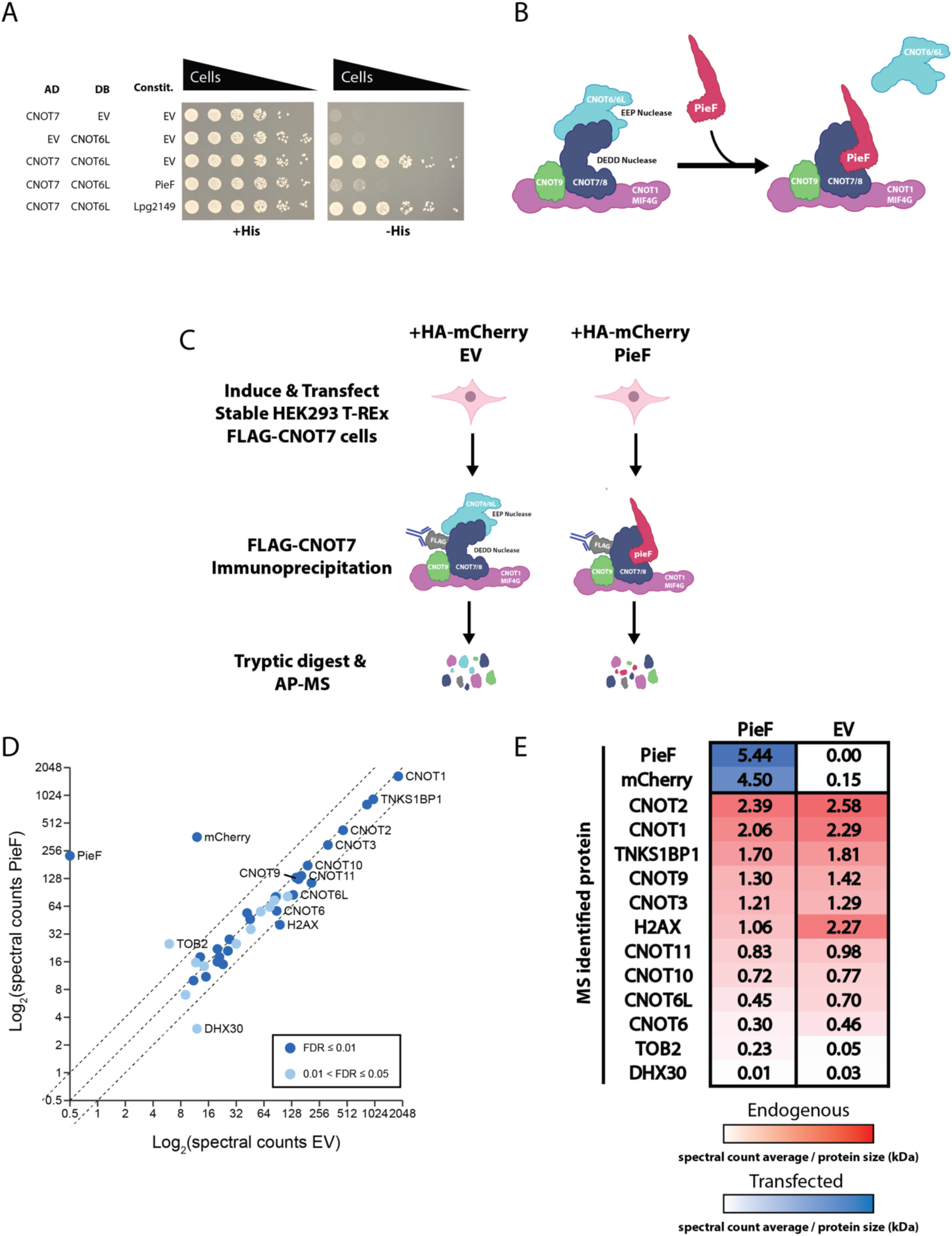
PieF impairs EEP nuclease incorporation into CCR4-NOT. **A)** Y2H assay with a constitutively expressed third protein demonstrates that PieF specifically disrupts the Y2H signal between DEDD nuclease subunit CNOT7 and EEP nuclease subunit CNOT6L. Y2H spot-dilution was performed with undiluted culture (leftmost column #1) as well as serial 5-fold dilutions of culture from OD600=1.0 to 0.0016 (columns 2-6). **B)** Schematic model for EEP nuclease displacement by PieF in a binding site overlap model. **C)** FLAG AP-MS schematic. A stable Flp-In™ T-REx™ 293 cell line with FLAG-tagged CNOT7 is transfected with HA-mCherry vectors and subjected to FLAG AP-MS to assess subunit composition changes in CCR4-NOT. **D)** A prey gene spectral abundance scatter plot comparison depicting SAINT analysis of AP-MS results. The plot illustrates the depletion of the EEP nuclease module relative to when an empty vector is transfected. Canonical CCR4-NOT components and TNKS1BP1 are labelled, as well as prey that exceeds the 2-fold change thresholds (dashed lines). **E)** Heatmap representation for average spectral counts normalized to protein size between PieF and EV treatment.

To further support the observation that PieF may impair CNOT6/6L recruitment to the CCR4-NOT complex, we immunoprecipitated FLAG-tagged CNOT7 (expressed from stable cell pools via a tetracycline-inducible promoter) from cells transiently transfected with either HA-mCherry tagged PieF, or HA-mCherry vector alone (Fig. 4C). Following PieF expression, we observed a decrease of 36% of CNOT6 and 35% of CNOT6L spectral counts co-precipitated by CNOT7 relative to empty vector (Fig. 4D & 4E), corroborating Y2H results. Interestingly other canonical components depicted did not show such marked changes in abundance between treatment conditions, suggesting that the canonical CCR4-NOT complex remains largely intact. CNOT11 was the next most differentially enriched, with 84% of total peptides being captured in the PieF co-precipitation relative to empty vector. Collectively our yeast two-hybrid, and mass spectrometry results suggest that PieF alters the subunit composition of CCR4-NOT, specifically by influencing EEP nuclease (CNOT6/6L) incorporation.

### Overexpression of PieF in mammalian cells results in the nuclear exclusion of CNOT7

Given that CNOT7 can shuttle between cellular compartments during the cell cycle (126), we also assayed whether PieF influenced the localization of CNOT7 following ectopic expression in HEK293T cells. The catalytic subunit was tagged with eGFP at its N-terminus and localization was monitored by fluorescence microscopy in the presence of HA-mCherry tagged PieF, or Lpg2149. Most cells in the HA-mCherry vector, or HA-mCherry Lpg2149 treatment demonstrated diffuse pan cellular localization of eGFP-CNOT7 (Fig. 5A), consistent with other published CNOT7 localization data (126). In contrast, HA-mCherry PieF expression resulted in a significant proportion of the cell population showing a diminished eGFP signal within the nucleus (Fig.s 5A and B). PieF appeared to localize throughout the cell, analogous to TOB proteins (127). Taken together, our data demonstrate that PieF can redirect the DEDD nuclease from the nucleus.

**Figure 5.**
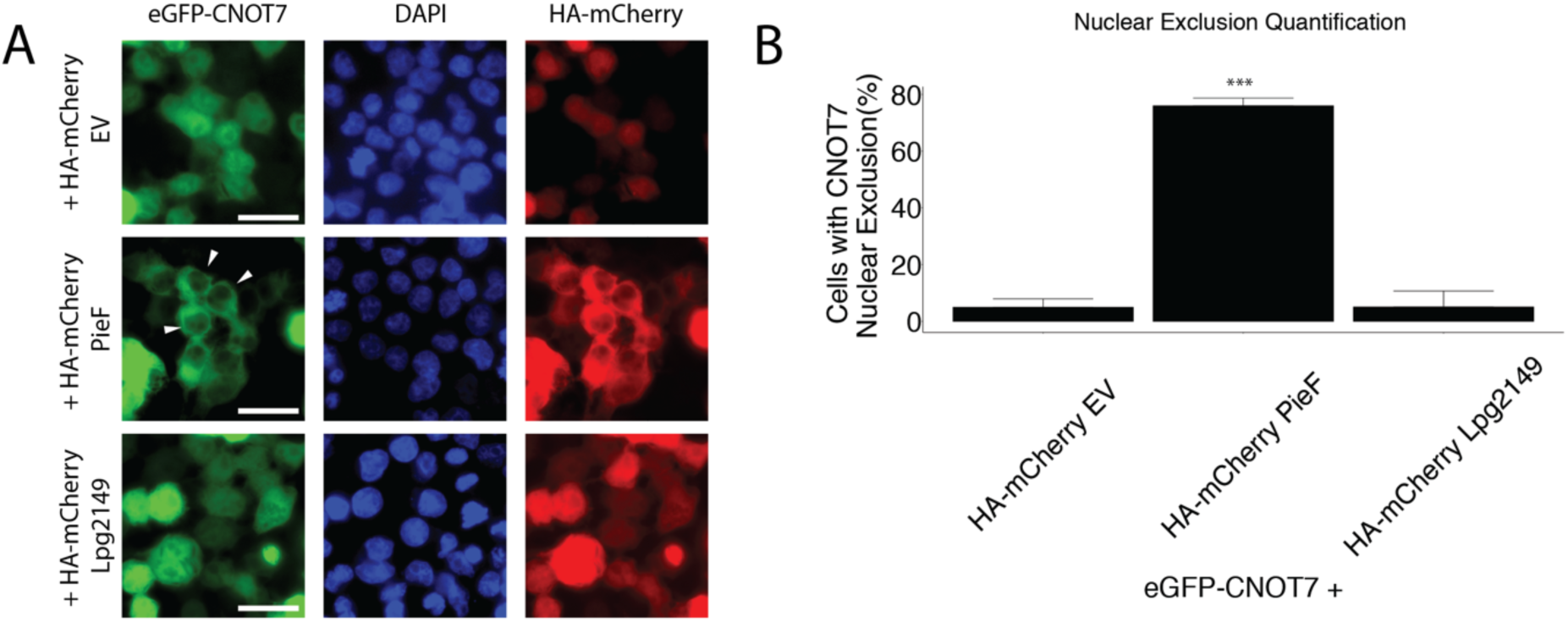
PieF restricts CNOT7 nuclear localization following ectopic expression in HEK293T cells. **A)** Transient transfection of HEK293T with eGFP-CNOT7 and HA-mCherry PieF causes a nuclear exclusion phenotype that is specific to PieF and visible under a 40X oil-immersion objective. The scale bar represents 50 μm. Arrows illustrate representative nuclear exclusion phenotypes. 4’, 6- diamidino-2-phenylindole (DAPI) illustrates nuclear staining. Images are representative of at least two biological replicates. **B)** Manual quantification of cells (>100 per replicate) with nuclear-excluded eGFP-CNOT7 shows a statistically significant difference in the PieF populations relative to other treatments using a Student’s t-test (*** = p < 0.01) across two biological replicates.

### Overexpression of PieF in yeast phenocopies deletion of the CNOT7/8 ortholog, *Pop2*

CCR4-NOT was first identified and studied in *S. cerevisiae* (93). In yeast, deletion of the ortholog of *CNOT7/8*, called *POP2,* leads to several cellular phenotypes, including reduced growth rate in rich medium, and sensitivity to the transcriptional elongation inhibitor 6-azauracil (128). CCR4- NOT components have synthetic lethal genetic interactions with transcriptional elongation factors and can stimulate the elongation of arrested RNA polymerase II *in vitro* supporting a role in transcriptional elongation (128–131). 6-azauracil depletes cellular pools of GTP by inhibiting IMP-dehydrogenase, which also blocks transcriptional elongation by RNA polymerase II (132). To determine if PieF could re-direct CCR4-NOT activity across diverse eukaryotic organisms we assayed the growth of *S. cerevisiae* strains expressing PieF or lacking *POP2* in the presence of 6-azauracil. Consistent with our previous data suggesting CCR4-NOT targeting by PieF, expression of PieF in *S. cerevisiae* resulted in enhanced susceptibility to 6-azauracil (Fig. 6A), phenocopying the deletion of *POP2*. PieF expression also resulted in a modest, but insignificant growth inhibition in the absence of 6-azauracil, which is similar, although not as severe, as deletion of *POP2* (Fig. 6B). This phenotype was not observed when a negative control (the unrelated effector Lpg2149) was expressed. Taken together, PieF expression phenocopies the deletion of *POP2* in *S. cerevisiae*, suggesting that PieF impinges upon CCR4-NOT activity across diverse eukaryotic species.

**Figure 6.**
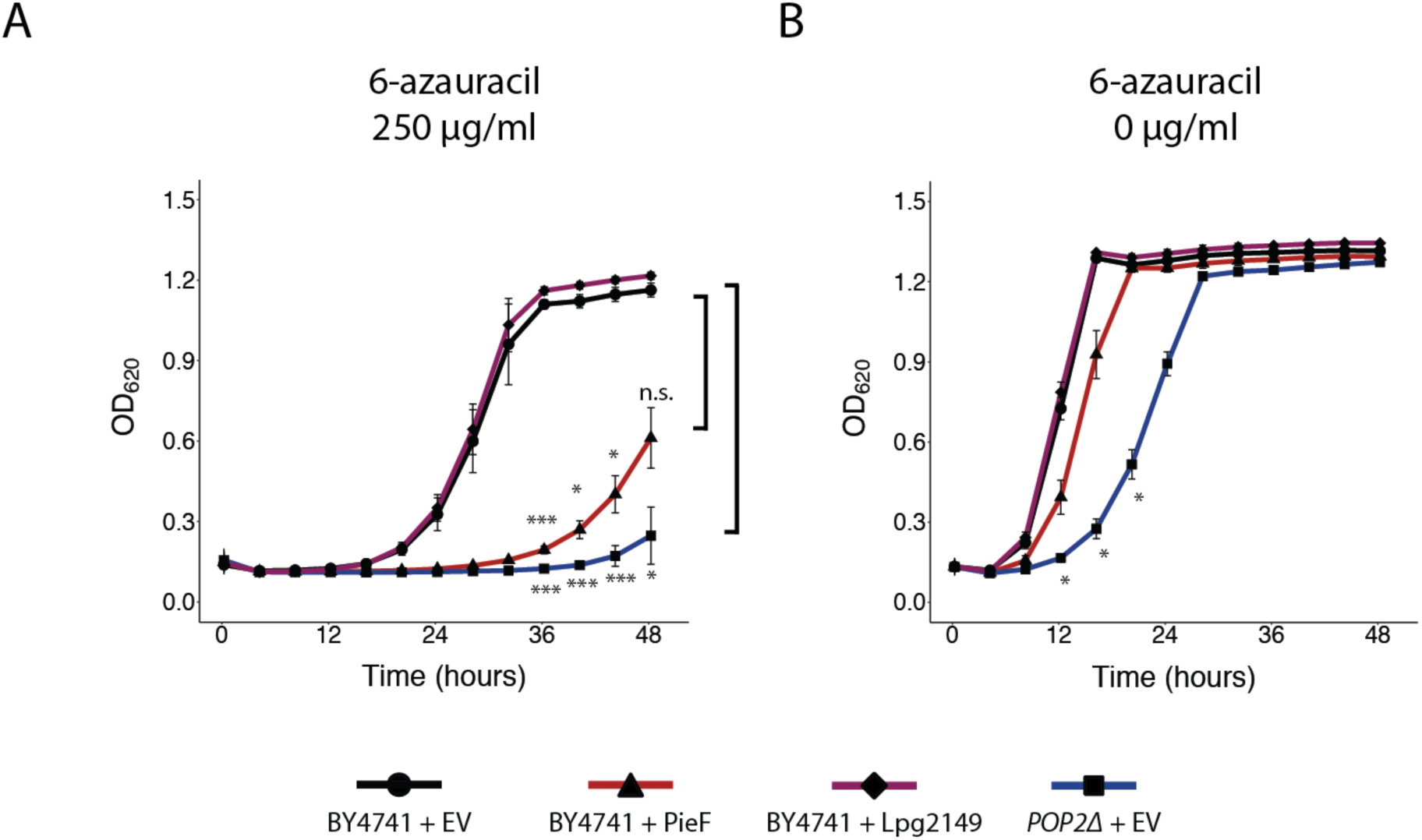
PieF modulates CCR4-NOT in yeast. **A)** Growth curves of wild-type BY4741 or BY4741 *pop2Δ* carrying overexpression vectors of PieF, Lpg2149 or empty vector control (EV) in the presence of 250 μg/ml 6-azauracil. The average of two experiments and the standard error of the mean is plotted. * p < 0.05, *** p < 0.01 by Benjamini-Hochberg corrected Students t-test. Deletion of the yeast *CNOT7* ortholog *POP2* leads to a growth defect in the presence of 6-azauracil as does expression of PieF, but not with the expression of Lpg2149 or an EV control. **B)** PieF expression as above also demonstrates a modest but insignificant growth defect in the absence of 6-azauracil.

## DISCUSSION

The work presented here reveals a new function for another effector within the *Pie* locus. While PieA, PieE, and PieG appear to influence vesicular trafficking and LCV motility, PieF appears to instead target distinct eukaryotic biology. To our knowledge, PieF is the first described virulence protein encoded by a human bacterial pathogen that intersects the eukaryotic CCR4-NOT deadenylase machinery. The CCR4-NOT complex is found across eukaryotes (54), making it an attractive target for a bacterial pathogen that can replicate in host cells as evolutionarily diverse as protists and human macrophages (5). Despite having conserved CCR4-NOT components, the protozoan CCR4-NOT complex is largely unstudied (133, 134). CCR4-NOT components (including CNOT7) are found in protists such as *Entamoeba histolytica* (133, 134). Furthermore, in some trypanosome species, CNOT7 is the only conserved deadenylase and essential for growth (133–137). The DEDD nucleases (CNOT7/8) are considered the primordial nuclease module of CCR4-NOT, being much more conserved throughout eukaryotic evolution than the EEP nuclease module (136). As a pathogenic target CNOT7/8 is much more likely to be conserved across evolutionarily diverse eukaryotic hosts and represents a prime pathogenic target. Indeed, our *S. cerevisiae* data suggests that PieF can redirect CCR4-NOT activity across evolutionarily distant eukaryotic species.

In the literature, there is a prominent example of a pathogenic factor that targets the CNOT7 subunit of CCR4-NOT (102). The *Xanthomonas citri* effector, PthA4, inhibits *Citrus sinensis* CNOT7 activity *in vitro* and promotes transcription and translation of factors that facilitate *X. citri* canker formation (102). This is one of several examples of the CCR4-NOT complex providing protective effects against environmental stressors and pathogens in plants (102, 138–140). There are also studies demonstrating that CNOT7 is required for the development of gametocytes in the malarial parasite *Plasmodium falciparum* (141), and CNOT7 orthologs also play a role in diverse stress response phenotypes within model and pathogenic fungi alike (142). As a central regulator of eukaryotic gene expression, with roles in immune signalling (143) and stress response pathways (142), there are several ways in which modulation of the CCR4-NOT complex by PieF could influence host cell gene expression ranging from generalized changes to mRNA stability to the modulation of specific targets. Generalized transcriptomic changes may occur if PieF can recruit CCR4-NOT to mRNA without clear specificity like the BTG/TOB proteins (96, 144). Alternatively, PieF could direct CNOT7 to specific mRNA transcripts through association with RNA-binding proteins (e.g., TTP, Nanos, Pumilio, PABPC1), or the miRNA machinery (TNRC6A, TNRC6B, TNRC6C) which can selectively recruit CCR4-NOT for mRNA silencing and deadenylation (54). Our observation of PieF- directed CNOT7 localization changes is perhaps informative in this regard, suggesting an upstream influence on CNOT7 activity that may nevertheless modulate specific classes of transcripts based on one or more general characteristics.

In the environment, resources are often scarce, and many protozoan cells are in stationary phase (145). Endogenous CNOT7 is almost exclusively nuclear in starved mammalian MRC5 cells (126), and this may best approximate the expected configuration of amoebal CCR4-NOT in a nutrient-poor environment as well. PieF expression appears to favour cytoplasmic CNOT7. During *L. pneumophila* infection of one or more protozoan hosts, PieF may re-direct nuclear CNOT7 activity towards the cytoplasm, which might ultimately favour bacterial replication by destabilizing specific mRNA subsets, blocking the host cell cycle, or facilitating translational repression through CCR4-NOT recruitment. It has been shown that cytoplasmic localization of TOB was critical for its ability to inhibit cellular proliferation (127). *L. pneumophila* has been previously shown to actively arrest the host cell cycle in both mammalian, and protozoan cells through the activity of numerous effectors (47, 145–147), aiding the formation of a replication permissive compartment. During infection, it is imperative for the bacterium to block host entry into S-phase, as this cell cycle stage destabilizes the LCV and prevents bacterial replication (145). The highly conserved CCR4-NOT complex presents an enticing and unexplored avenue for modulating the host from a pathogenic perspective (91, 109, 110).

Within this context, the similarities between PieF and the BTG/TOB family are striking. Like PieF, BTG/TOB proteins interact directly with CNOT7/8 of the CCR4-NOT complex (91, 148, 149). PieF (Fig. 2A), BTG2 (107) and TOB (108) have been shown to directly impair CNOT7 catalytic activity *in vitro*. Conversely, tethering of each component within cells facilitates mRNA repression and reduction in steady-state levels (68, 113, 118) (Fig. 2B). These seemingly contradictory results may be explained by the requirement of an accessory protein to mediate PieF-bound CCR4-NOT recruitment to mRNA. Previous studies have shown that the addition of PABP to *in vitro* reactions of CNOT7-TOB can stimulate nuclease activity (72). More recent work has shown that PABP occupancy may influence the catalytic activity of each CCR4-NOT deadenylase module (67). BTG/TOB proteins are antiproliferative and can halt mammalian cell-cycle progression when overexpressed, in a manner that is dependent on interaction with CCR4-NOT (91, 109, 110, 113). If PieF mimics BTG/TOB protein molecular function its role may be to abruptly shut down the host cell cycle through antiproliferative effects (91, 109, 110), creating a replication permissive environment in the host (145).

In the future, it will be important to delineate which mRNA transcripts are influenced by PieF both ectopically, and during infection. Bulk RNA-sequencing following transfection of PieF might reveal specific transcripts that are sensitive to its modulation of CNOT7. Additionally, global changes to poly(A) tail length could be monitored using a TAIL-seq-based approach (150) to see whether PieF can trigger general mRNA deadenylation (96). If the role of PieF is to block host cell-cycle progression, the target transcripts should largely overlap with the BTG/TOB family (96). Such a role would explain why there is no replication defect of a PieF mutant in terminally differentiated macrophages which are not progressing through the cell cycle (Fig. S1B). It is worth noting however that a recent systematic mutagenesis study did not identify an essential role of PieF in four natural protozoan hosts (89). In this light, it will be important to explore potential genetic redundancy between PieF and other effectors (24, 26). Indeed, the conservation of CCR4-NOT and the influence of the complex on so many facets of fundamental eukaryotic biology make it likely that additional effectors interface with this pathway.

## AVAILABILITY

CCR4-NOT subunit composition data has been deposited as a complete submission to the MassIVE repository (https://massive.ucsd.edu/ProteoSAFe/static/massive.jsp)

## ACCESSION NUMBERS

MassIVE repository accession number: MSV000089325 ProteomeXchange accession: PXD033482

## ACKNOWLEDGEMENT

The authors thank Dylan Valleau, Alexei Savchenko, members of the Savchenko laboratory, and the BioZone mass-spectrometry facility for their help with cloning, protein purification, and AP-MS identification of PieF host targets. We also thank Maciej Jagielnicki and Trevor Moraes for assistance with BLI. Additionally, we would like to thank Marc Fabian for providing tethering constructs and experimental consultation, Karen Maxwell for providing bacterial two-hybrid constructs, and Dae-Kyum Kim and Fritz Roth for providing human cDNA clones, and experimental advice. Finally, we thank members of the Ensminger laboratory for their suggestions and careful reading of the manuscript.

---

**Figure S1 PieF is a non-essential effector in the *pie* locus. A)** *PieF* (Lpg1972) is found within the *pie* locus of *L. pneumophila* (Ninio *et al*., 2009). The innermost ring of the Circos plot represents the distribution of effectors across the *L. pneumophila* Str. Philadelphia-1 genome as a barcode. The *pie* locus is emphasized, and *pie* effectors are colored purple. The outermost ring is a bar plot representation of effector conservation across 41 sequenced *Legionella* species (Burstein *et al*., 2016). **B)** Intracellular replication was measured by CFU plating at three time points (2, 24, and 48 h) during U937 infection (initial MOI 0.05) onto charcoal yeast. Deletion of *pieF* shows no replication defect relative to wildtype, the *dotA*- mutant lacking a functional type IV secretion system fails to replicate intracellularly.

**Table S1. Plasmids used in this study.**

**Table S2. Bacterial Strains used in this study.**

**Table S3. *L. pneumophila* strains used in this study.**

**Table S4. Primers & Oligonucleotide(s) used in this study**

**Table S5. Yeast strains used in this study.**

**Table S6. Mammalian cell lines used in this study**

**Table S7. Proteins removed by ProHits artifact, keratin, and ribosomal biofilters.**

## REFERENCES

1. Fliermans, C.B., Orrison, L.H., Smith, J., Cherry, W.B., Tison, D.L. and Pope, D.H. (2005) Ecological Distribution of *Legionella pneumophila*. Appl. Environ. Microbiol.

2. Morris, G.K., Patton, C.M., Feeley, J.C., Johnson, S.E., Gorman, G., Skaliy, P., Mallison, G.F., Poloto, B.D. and Mackel, D.C. (1979) Isolation of the Legionnaires’ Disease Bacterium from Environmental Samples. Annals of Internal Medicine, 90, 664–666.

3. Rowbotham, T.J. (1980) Preliminary report on the pathogenicity of *Legionella pneumophila* for freshwater and soil amoebae. J. Clin. Pathol., 33, 1179–1183.

4. Fields, B.S. (1996) The molecular ecology of *Legionellae*. Trends in Microbiology, 4, 286–290.

5. Boamah, D.K., Zhou, G., Ensminger, A.W. and O’Connor, T.J. (2017) From Many Hosts, One Accidental Pathogen: The Diverse Protozoan Hosts of *Legionella*. Front. Cell. Infect. Microbiol., 7, 21660–17.

6. Rowbotham, T.J. (1983) Isolation of *Legionella pneumophila* from clinical specimens via amoebae, and the interaction of those and other isolates with amoebae. J. Clin. Pathol., 36, 978–986.

7. Thomas, V., Bouchez, T., Nicolas, V., Robert, S., Loret, J.F. and Lévi, Y. (2004) Amoebae in domestic water systems: resistance to disinfection treatments and implication in *Legionella* persistence. J Appl Microbiol, 97, 950–963.

8. Ikedo, M. and Yabuuchi, E. (1986) Ecological Studies of *Legionella* Species I. Viable Counts of *Legionella pneumophila* in Cooling Tower Water. Microbiol. Immunol., 30, 413–423.

9. Breiman, R.F., Fields, B.S., Sanden, G.N., Volmer, L., Meier, A. and Spika, J.S. (1990) Association of Shower Use With Legionnaires’ Disease: Possible Role of Amoebae. JAMA, 263, 2924–2926.

10. Yamamoto, H., Sugiura, M., Shinji, K., Takayuki, E., Ikedo, M. and Yabuuchi, E. (1992) Factors Stimulating Propagation of *Legionellae* in Cooling Tower. Appl. Environ. Microbiol.

11. Fields, B.S., Benson, R.F. and Besser, R.E. (2002) *Legionella* and Legionnaires’ disease: 25 years of investigation. Clin. Microbiol. Rev., 15, 506–526.

12. Lasheras, A., Boulestreau, H., Rogues, A.-M., Ohayon-Courtes, C., Labadie, J.-C. and Gachie, J.-P. (2006) Influence of amoebae and physical and chemical characteristics of water on presence and proliferation of *Legionella* species in hospital water systems. Am J Infect Control, 34, 520–525.

13. Brousseau, N., Lévesque, B., Guillemet, T.A., Cantin, P., Gauvin, D., Giroux, J.-P., Gingras, S., Proulx, F., Côté, P.-A. and Dewailly, E. (2013) Contamination of public whirlpool spas: factors associated with the presence of *Legionella* spp., *Pseudomonas aeruginosa* and *Escherichia coli*. Int J Environ Health Res, 23, 1–15.

14. Thomas, J.M., Thomas, T., Stuetz, R.M. and Ashbolt, N.J. (2014) Your garden hose: a potential health risk due to *Legionella* spp. growth facilitated by free-living amoebae. Environ Sci Technol, 48, 10456–10464.

15. Horwitz, M.A. (1983) Formation of a novel phagosome by the Legionnaires’ disease bacterium (*Legionella pneumophila*) in human monocytes. J. Exp. Med., 158, 1319–1331.

16. Horwitz, M.A. (1983) The Legionnaires’ disease bacterium (*Legionella pneumophila*) inhibits phagosome-lysosome fusion in human monocytes. J. Exp. Med., 158, 2108–2126.

17. Horwitz, M.A. and Maxfield, F.R. (1984) *Legionella pneumophila* inhibits acidification of its phagosome in human monocytes. J Cell Biol, 99, 1936–1943.

18. Bozue, J.A. and Johnson, W. (1996) Interaction of *Legionella pneumophila* with *Acanthamoeba castellanii*: uptake by coiling phagocytosis and inhibition of phagosome-lysosome fusion. Infection and Immunity, 64, 668–673.

19. Nagai, H., Kagan, J.C., Zhu, X., Kahn, R.A. and Roy, C.R. (2002) A bacterial guanine nucleotide exchange factor activates ARF on *Legionella* phagosomes. Science (New York, N.Y.), 295, 679– 682.

20. Conover, G.M., Derre, I., Vogel, J.P. and Isberg, R.R. (2003) The *Legionella pneumophila* LidA protein: a translocated substrate of the Dot/Icm system associated with maintenance of bacterial integrity. Molecular Microbiology, 48, 305–321.

21. Chen, J., de Felipe, K.S., Clarke, M., Lu, H., Anderson, O.R., Segal, G. and Shuman, H.A. (2004) *Legionella* Effectors That Promote Nonlytic Release from Protozoa. Science, 303, 1358–1361.

22. Luo, Z.Q. and Isberg, R.R. (2004) Multiple substrates of the *Legionella pneumophila* Dot/Icm system identified by interbacterial protein transfer. PNAS, 101, 841–846.

23. Ensminger, A.W. (2016) *Legionella pneumophila*, armed to the hilt: justifying the largest arsenal of effectors in the bacterial world. Curr. Opin. Microbiol., 29, 74–80.

24. O’Connor, T.J., Adepoju, Y., Boyd, D. and Isberg, R.R. (2011) Minimization of the *Legionella pneumophila* genome reveals chromosomal regions involved in host range expansion. Proceedings of the National Academy of Sciences of the United States of America, 108, 14733– 14740.

25. O’Connor, T.J., Boyd, D., Dorer, M.S. and Isberg, R.R. (2012) Aggravating Genetic Interactions Allow a Solution to Redundancy in a Bacterial Pathogen. Science, 338, 1440–1444.

26. Ghosh, S. and O’Connor, T.J. (2017) Beyond Paralogs: The Multiple Layers of Redundancy in Bacterial Pathogenesis. Front. Cell. Infect. Microbiol., 7, 467.

27. Chien, M., Morozova, I., Shi, S., Sheng, H., Chen, J., Gomez, S.M., Asamani, G., Hill, K., Nuara, J., Feder, M., et al. (2004) The genomic sequence of the accidental pathogen *Legionella pneumophila*. Science (New York, N.Y.), 305, 1966–1968.

28. Burstein, D., Amaro, F., Zusman, T., Lifshitz, Z., Cohen, O., Gilbert, J.A., Pupko, T., Shuman, H.A. and Segal, G. (2016) Genomic analysis of 38 *Legionella* species identifies large and diverse effector repertoires. Nat Genet, 48, 167–175.

29. Gomez-Valero, L., Rusniok, C., Carson, D., Mondino, S., Pérez-Cobas, A.E., Rolando, M., Pasricha, S., Reuter, S., Demirtas, J., Crumbach, J., et al. (2019) More than 18, 000 effectors in the *Legionella* genus genome provide multiple, independent combinations for replication in human cells. Proceedings of the National Academy of Sciences of the United States of America, 116, 2265–2273.

30. Ninio, S., Celli, J. and Roy, C.R. (2009) A *Legionella pneumophila* Effector Protein Encoded in a Region of Genomic Plasticity Binds to Dot/Icm-Modified Vacuoles. PLoS Pathog, 5, e1000278.

31. Mousnier, A., Schroeder, G.N., Stoneham, C.A., So, E.C., Garnett, J.A., Yu, L., Matthews, S.J., Choudhary, J.S., Hartland, E.L. and Frankel, G. (2014) A New Method To Determine *In Vivo* Interactomes Reveals Binding of the *Legionella pneumophila* Effector PieE to Multiple Rab GTPases. mBio, 5, e01148–14–e01148–14.

32. Rothmeier, E., Pfaffinger, G., Hoffmann, C., Harrison, C.F., Grabmayr, H., Repnik, U., Hannemann, M., Wölke, S., Bausch, A., Griffiths, G., et al. (2013) Activation of Ran GTPase by a *Legionella* Effector Promotes Microtubule Polymerization, Pathogen Vacuole Motility and Infection. PLoS Pathog, 9, e1003598–17.

33. de Felipe, K.S., Pampou, S., Jovanovic, O.S., Pericone, C.D., Ye, S.F., Kalachikov, S. and Shuman, H.A. (2005) Evidence for Acquisition of *Legionella* Type IV Secretion Substrates via Interdomain Horizontal Gene Transfer. Journal of Bacteriology, 187, 7716–7726.

34. de Felipe, K.S., Glover, R.T., Charpentier, X., Anderson, O.R., Reyes, M., Pericone, C.D. and Shuman, H.A. (2008) *Legionella* Eukaryotic-Like Type IV Substrates Interfere with Organelle Trafficking. PLoS Pathog, 4, e1000117.

35. Ivanov, S.S., Charron, G., Hang, H.C. and Roy, C.R. (2010) Lipidation by the Host Prenyltransferase Machinery Facilitates Membrane Localization of *Legionella pneumophila* Effector Proteins. J. Biol. Chem., 285, 34686–34698.

36. Cornejo, E., Schlaermann, P. and Mukherjee, S. (2021) How to rewire the host cell: A home improvement guide for intracellular bacteria. 10.1083/jcb.201701095.

37. Hempstead, A.D. and Isberg, R.R. (2015) Inhibition of host cell translation elongation by *Legionella pneumophila* blocks the host cell unfolded protein response. PNAS, 112, E6790–E6797.

38. Shin, S., Case, C.L., Archer, K.A., Nogueira, C.V., Kobayashi, K.S., Flavell, R.A., Roy, C.R. and Zamboni, D.S. (2008) Type IV Secretion-Dependent Activation of Host MAP Kinases Induces an Increased Proinflammatory Cytokine Response to *Legionella pneumophila*. PLoS Pathog, 4, e1000220–13.

39. Fontana, M.F., Banga, S., Barry, K.C., Shen, X., Tan, Y., Luo, Z.-Q. and Vance, R.E. (2011) Secreted Bacterial Effectors That Inhibit Host Protein Synthesis Are Critical for Induction of the Innate Immune Response to Virulent *Legionella pneumophila*. PLoS Pathog, 7, e1001289.

40. Fontana, M.F., Shin, S. and Vance, R.E. (2012) Activation of host mitogen-activated protein kinases by secreted *Legionella pneumophila* effectors that inhibit host protein translation. Infection and Immunity, 80, 3570–3575.

41. Barry, K.C., Fontana, M.F., Portman, J.L., Dugan, A.S. and Vance, R.E. (2013) IL-1α signaling initiates the inflammatory response to virulent *Legionella pneumophila in vivo*. J. Immunol., 190, 6329–6339.

42. Asrat, S., Dugan, A.S. and Isberg, R.R. (2014) The Frustrated Host Response to *Legionella pneumophila* Is Bypassed by MyD88-Dependent Translation of Pro-inflammatory Cytokines. PLoS Pathog, 10, e1004229–15.

43. Samara, H.A., Richards, A.M., Price, C.T.D., Dwingelo, Von, J.E. and Abu Kwaik, Y. (2021) Amoeba host-*Legionella* synchronization of amino acid auxotrophy and its role in bacterial adaptation and pathogenic evolution. Environmental Microbiology, 16, 350–358.

44. Belyi, Y. (2020) Targeting Eukaryotic mRNA Translation by *Legionella pneumophila*. Frontiers in Molecular Biosciences, 10.3389/fmolb.2020.00080.

45. De Leon, J.A., Qiu, J., Nicolai, C.J., Counihan, J.L., Barry, K.C., Xu, L., Lawrence, R.E., Castellano, B.M., Zoncu, R., Nomura, D.K., et al. (2017) Positive and Negative Regulation of the Master Metabolic Regulator mTORC1 by Two Families of *Legionella pneumophila* Effectors. Cell Reports, 21, 2031–2038.

46. Denzer, L., Schroten, H. and Schwerk, C. (2020) From Gene to Protein—How Bacterial Virulence Factors Manipulate Host Gene Expression During Infection. International Journal of Molecular Sciences, 10.3390/ijms21103730.

47. Sol, A., Lipo, E., de Jesús-Díaz, D.A., Murphy, C., Devereux, M. and Isberg, R.R. (2019) *Legionella pneumophila* translocated translation inhibitors are required for bacterial-induced host cell cycle arrest. Proceedings of the National Academy of Sciences of the United States of America, 116, 3221–3228.

48. Belyi, I., Popoff, M.R. and Cianciotto, N.P. (2003) Purification and characterization of a UDP-glucosyltransferase produced by *Legionella pneumophila*. Infection and Immunity, 71, 181–186.

49. Belyi, Y., Niggeweg, R., Opitz, B., Vogelsgesang, M., Hippenstiel, S., Wilm, M. and Aktories, K. (2006) *Legionella pneumophila* glucosyltransferase inhibits host elongation factor 1A. PNAS, 103, 16953–16958.

50. Shen, X., Banga, S., Liu, Y., Xu, L., Gao, P., Shamovsky, I., Nudler, E. and Luo, Z.-Q. (2009) Targeting eEF1A by a *Legionella pneumophila* effector leads to inhibition of protein synthesis and induction of host stress response. Cellular Microbiology, 11, 911–926.

51. Joseph, A.M., Pohl, A.E., Ball, T.J., Abram, T.G., Johnson, D.K., Geisbrecht, B.V. and Shames, S.R. (2020) The *Legionella pneumophila* Metaeffector Lpg2505 (MesI) Regulates SidI-Mediated Translation Inhibition and Novel Glycosyl Hydrolase Activity. Infection and Immunity, 88, 1–17.

52. Flayhan, A., Bergé, C., Baïlo, N., Doublet, P., Bayliss, R. and Terradot, L. (2015) The structure of *Legionella pneumophila* LegK4 type four secretion system (T4SS) effector reveals a novel dimeric eukaryotic-like kinase. Sci. Rep., 5, 14602.

53. Moss, S.M., Taylor, I.R., Ruggero, D., Gestwicki, J.E., Shokat, K.M. and Mukherjee, S. (2019) A *Legionella pneumophila* Kinase Phosphorylates the Hsp70 Chaperone Family to Inhibit Eukaryotic Protein Synthesis. Cell Host Microbe, 25, 454–462.e6.

54. Chalabi Hagkarim, N. and Grand, R.J. (2020) The Regulatory Properties of the Ccr4-Not Complex. Cells, 9, 2379.

55. Villanyi, Z. and Collart, M.A. (2015) Ccr4-Not is at the core of the eukaryotic gene expression circuitry. Biochem Soc Trans, 43, 1253–1258.

56. Collart, M.A. and Panasenko, O.O. (2017) The Ccr4-Not Complex: Architecture and Structural Insights. Subcell Biochem, 83, 349–379.

57. Collart, M.A. (2016) The Ccr4-Not complex is a key regulator of eukaryotic gene expression. WIREs RNA, 7, 438–454.

58. Mostafa, D., Takahashi, A., Yanagiya, A., Yamaguchi, T., Abe, T., Kureha, T., Kuba, K., Kanegae, Y., Furuta, Y., Yamamoto, T., et al. (2020) Essential functions of the CNOT7/8 catalytic subunits of the CCR4-NOT complex in mRNA regulation and cell viability. RNA Biology, 17, 403–416.

59. Petit, A.-P., Wohlbold, L., Bawankar, P., Huntzinger, E., Schmidt, S., Izaurralde, E. and Weichenrieder, O. (2012) The structural basis for the interaction between the CAF1 nuclease and the NOT1 scaffold of the human CCR4–NOT deadenylase complex. Nucleic Acids Research, 40, 11058–11072.

60. Basquin, J., Roudko, V.V., Rode, M., Basquin, C., Séraphin, B. and Conti, E. (2012) Architecture of the Nuclease Module of the Yeast Ccr4-Not Complex: the Not1-Caf1-Ccr4 Interaction. Molecular Cell, 48, 207–218.

61. Lau, N.-C., Kolkman, A., van Schaik, F.M.A., Mulder, K.W., Pijnappel, W.W.M.P., Heck, A.J.R. and Timmers, H.T.M. (2009) Human Ccr4-Not complexes contain variable deadenylase subunits. Biochem J, 422, 443–453.

62. Morita, M., Suzuki, T., Nakamura, T., Yokoyama, K., Miyasaka, T. and Yamamoto, T. (2007) Depletion of mammalian CCR4b deadenylase triggers elevation of the p27Kip1 mRNA level and impairs cell growth. Mol. Cell. Biol., 27, 4980–4990.

63. Yamashita, A., Chang, T.-C., Yamashita, Y., Zhu, W., Zhong, Z., Chen, C.-Y.A. and Shyu, A.-B. (2005) Concerted action of poly(A) nucleases and decapping enzyme in mammalian mRNA turnover. Nature Structural & Molecular Biology, 12, 1054–1063.

64. Liu, H.Y., Badarinarayana, V., Audino, D.C., Rappsilber, J., Mann, M. and Denis, C.L. (1998) The NOT proteins are part of the CCR4 transcriptional complex and affect gene expression both positively and negatively. EMBO J, 17, 1096–1106.

65. Draper, M.P., Salvadore, C. and Denis, C.L. (1995) Identification of a mouse protein whose homolog in *Saccharomyces cerevisiae* is a component of the CCR4 transcriptional regulatory complex. Mol. Cell. Biol., 15, 3487–3495.

66. Mittal, S., Aslam, A., Doidge, R., Medica, R. and Winkler, G.S. (2011) The Ccr4a (CNOT6) and Ccr4b (CNOT6L) deadenylase subunits of the human Ccr4-Not complex contribute to the prevention of cell death and senescence. Molecular Biology of the Cell, 22, 748–758.

67. Yi, H., Park, J., Ha, M., Lim, J., Chang, H. and Kim, V.N. (2018) PABP Cooperates with the CCR4- NOT Complex to Promote mRNA Deadenylation and Block Precocious Decay. Molecular Cell, 70, 1081–1088.e5.

68. Mauxion, F., Faux, C. and Séraphin, B. (2008) The BTG2 protein is a general activator of mRNA deadenylation. The EMBO Journal, 27, 1039–1048.

69. Stupfler, B., Birck, C., Séraphin, B. and Mauxion, F. (2016) BTG2 bridges PABPC1 RNA-binding domains and CAF1 deadenylase to control cell proliferation. Nature Communications, 7, 10811.

70. Amine, H., Ripin, N., Sharma, S., Stoecklin, G., Allain, F.H., Séraphin, B. and Mauxion, F. (2021) A conserved motif in human BTG1 and BTG2 proteins mediates interaction with the poly(A) binding protein PABPC1 to stimulate mRNA deadenylation. RNA Biology, 18, 2450–2465.

71. Ezzeddine, N., Chang, T.-C., Zhu, W., Yamashita, A., Chen, C.-Y.A., Zhong, Z., Yamashita, Y., Zheng, D. and Shyu, A.-B. (2007) Human TOB, an antiproliferative transcription factor, is a poly(A)-binding protein-dependent positive regulator of cytoplasmic mRNA deadenylation. Mol. Cell. Biol., 27, 7791–7801.

72. Funakoshi, Y., Doi, Y., Hosoda, N., Uchida, N., Osawa, M., Shimada, I., Tsujimoto, M., Suzuki, T., Katada, T. and Hoshino, S.-I. (2007) Mechanism of mRNA deadenylation: evidence for a molecular interplay between translation termination factor eRF3 and mRNA deadenylases. Genes Dev., 21, 3135–3148.

73. Berger, K.H. and Isberg, R.R. (1993) Two distinct defects in intracellular growth complemented by a single genetic locus in *Legionella pneumophila*. Molecular Microbiology, 7, 7–19.

74. Ensminger, A.W., Yassin, Y., Miron, A. and Isberg, R.R. (2012) Experimental evolution of *Legionella pneumophila* in mouse macrophages leads to strains with altered determinants of environmental survival. PLoS Pathog, 8, e1002731.

75. Swanson, M.S. and Isberg, R.R. (1996) Identification of *Legionella pneumophila* mutants that have aberrant intracellular fates. Infection and Immunity, 64, 2585–2594.

76. Wu, B., Skarina, T., Yee, A., Jobin, M.-C., Dileo, R., Semesi, A., Fares, C., Lemak, A., Coombes, B.K., Arrowsmith, C.H., et al. (2010) NleG Type 3 effectors from enterohaemorrhagic *Escherichia coli* are U-Box E3 ubiquitin ligases. PLoS Pathog, 6, e1000960.

77. Quaile, A.T., Urbanus, M.L., Stogios, P.J., Nocek, B., Skarina, T., Ensminger, A.W. and Savchenko, A. (2015) Molecular Characterization of LubX: Functional Divergence of the U-Box Fold by *Legionella pneumophila*. Structure, 23, 1459–1469.

78. Liu, G., Zhang, J., Larsen, B., Stark, C., Breitkreutz, A., Lin, Z.-Y., Breitkreutz, B.-J., Ding, Y., Colwill, K., Pasculescu, A., et al. (2010) ProHits: integrated software for mass spectrometry-based interaction proteomics. Nat. Biotechnol., 28, 1015–1017.

79. Yu, H., Braun, P., Yildirim, M.A., Lemmens, I., Venkatesan, K., Sahalie, J., Hirozane-Kishikawa, T., Gebreab, F., Li, N., Simonis, N., et al. (2008) High-Quality Binary Protein Interaction Map of the Yeast Interactome Network. Science, 322, 104–110.

80. Hartley, J.L. (2000) DNA Cloning Using *In Vitro* Site-Specific Recombination. Genome Res., 10, 1788–1795.

81. Walhout, A.J., Temple, G.F., Brasch, M.A., Hartley, J.L., Lorson, M.A., van den Heuvel, S. and Vidal, M. (2000) GATEWAY recombinational cloning: application to the cloning of large numbers of open reading frames or ORFeomes. Methods Enzymol, 328, 575–592.

82. Yachie, N., Petsalaki, E., Mellor, J.C., Weile, J., Jacob, Y., Verby, M., Ozturk, S.B., Li, S., Cote, A.G., Mosca, R., et al. (2016) Pooled-matrix protein interaction screens using Barcode Fusion Genetics. Molecular Systems Biology, 12, 863–863.

83. Bickle, M.B.T., Dusserre, E., Moncorgé, O., Bottin, H. and Colas, P. (2006) Selection and characterization of large collections of peptide aptamers through optimized yeast two-hybrid procedures. Nature protocols, 1, 1066–1091.

84. Alberti, S., Gitler, A.D. and Lindquist, S. (2007) A suite of Gateway cloning vectors for high-throughput genetic analysis in *Saccharomyces cerevisiae*. Yeast, 24, 913–919.

85. Karimova, G., Pidoux, J., Ullmann, A. and Ladant, D. (1998) A bacterial two-hybrid system based on a reconstituted signal transduction pathway. PNAS, 95, 5752–5756.

86. Eng, J.K., Jahan, T.A. and Hoopmann, M.R. (2013) Comet: an open-source MS/MS sequence database search tool. PROTEOMICS, 13, 22–24.

87. Cox, J. and Mann, M. (2008) MaxQuant enables high peptide identification rates, individualized p.p.b.-range mass accuracies and proteome-wide protein quantification. Nat. Biotechnol., 26, 1367–1372.

88. Choi, H., Larsen, B., Lin, Z.-Y., Breitkreutz, A., Mellacheruvu, D., Fermin, D., Qin, Z.S., Tyers, M., Gingras, A.-C. and Nesvizhskii, A.I. (2011) SAINT: probabilistic scoring of affinity purification-mass spectrometry data. Nat Meth, 8, 70–73.

89. Park, J.M., Ghosh, S. and O’Connor, T.J. (2020) Combinatorial selection in amoebal hosts drives the evolution of the human pathogen *Legionella pneumophila*. Nature Microbiology, 5, 599–609.

90. Brown, C.E., Tarun, S.Z., Boeck, R. and Sachs, A.B. (1996) PAN3 encodes a subunit of the Pab1p- dependent poly(A) nuclease in *Saccharomyces cerevisiae*. Mol. Cell. Biol., 16, 5744–5753.

91. Ikematsu, N., Yoshida, Y., Kawamura-Tsuzuku, J., Ohsugi, M., Onda, M., Hirai, M., Fujimoto, J. and Yamamoto, T. (1999) Tob2, a novel anti-proliferative Tob/BTG1 family member, associates with a component of the CCR4 transcriptional regulatory complex capable of binding cyclin-dependent kinases. Oncogene, 18, 7432–7441.

92. Fields, S. and Song, O.-K. (1989) A novel genetic system to detect protein-protein interactions. Nature, 340, 245–246.

93. Denis, C.L. (1984) Identification of new genes involved in the regulation of yeast alcohol dehydrogenase II. Genetics, 108, 833–844.

94. Braun, P., Tasan, M., Dreze, M., Barrios-Rodiles, M., Lemmens, I., Yu, H., Sahalie, J.M., Murray, R.R., Roncari, L., De Smet, A.-S., et al. (2009) An experimentally derived confidence score for binary protein-protein interactions. Nat Meth, 6, 91–97.

95. Chen, J., Sawyer, N. and Regan, L. (2013) Protein-protein interactions: general trends in the relationship between binding affinity and interfacial buried surface area. Protein Sci, 22, 510–515.

96. Winkler, G.S. (2010) The mammalian anti-proliferative BTG/Tob protein family. J Cell Physiol, 222, 66–72.

97. Bai, Y., Tashiro, S., Nagatoishi, S., Suzuki, T., Yan, D., Liu, R., Tsumoto, K., Bartlam, M. and Yamamoto, T. (2015) Structural basis for inhibition of the Tob-CNOT7 interaction by a fragment screening approach. Protein Cell, 6, 924–928.

98. Ruan, L., Osawa, M., Hosoda, N., Imai, S., Machiyama, A., Katada, T., Hoshino, S.-I. and Shimada, I. (2010) Quantitative characterization of Tob interactions provides the thermodynamic basis for translation termination-coupled deadenylase regulation. J. Biol. Chem., 285, 27624–27631.

99. Viswanathan, P., Ohn, T., Chiang, Y.-C., Chen, J. and Denis, C.L. (2004) Mouse CAF1 Can Function As a Processive Deadenylase 3′–5′-Exonuclease *in vitro* but in Yeast the Deadenylase Function of CAF1 Is Not Required for mRNA Poly(A) Removal. Journal of Biological Chemistry, 279, 23988–23995.

100. Bianchin, C. (2005) Conservation of the deadenylase activity of proteins of the Caf1 family in human. RNA, 11, 487–494.

101. Zhang, Q., Yan, D., Guo, E., Ding, B., Yang, W., Liu, R., Yamamoto, T. and Bartlam, M. (2016) Structural basis for inhibition of the deadenylase activity of human CNOT6L. FEBS Letters, 590, 1270–1279.

102. Shimo, H.M., Terassi, C., Lima Silva, C.C., Zanella, J. de L., Mercaldi, G.F., Rocco, S.A. and Benedetti, C.E. (2019) Role of the *Citrus sinensis* RNA deadenylase CsCAF1 in citrus canker resistance. Mol Plant Pathol, 20, 1105–1118.

103. Meijer, H.A., Schmidt, T., Gillen, S.L., Langlais, C., Jukes-Jones, R., de Moor, C.H., Cain, K., Wilczynska, A. and Bushell, M. (2019) DEAD-box helicase eIF4A2 inhibits CNOT7 deadenylation activity. Nucleic Acids Research, 47, 8224–8238.

104. Horiuchi, M., Takeuchi, K., Noda, N., Muroya, N., Suzuki, T., Nakamura, T., Kawamura-Tsuzuku, J., Takahasi, K., Yamamoto, T. and Inagaki, F. (2009) Structural basis for the antiproliferative activity of the Tob-hCaf1 complex. Journal of Biological Chemistry, 284, 13244–13255.

105. Valleau, D., Quaile, A.T., Cui, H., Xu, X., Evdokimova, E., Chang, C., Cuff, M.E., Urbanus, M.L., Houliston, S., Arrowsmith, C.H., et al. (2018) Discovery of Ubiquitin Deamidases in the Pathogenic Arsenal of *Legionella pneumophila*. Cell Reports, 23, 568–583.

106. Wu, M., Reuter, M., Lilie, H., Liu, Y., Wahle, E. and Song, H. (2005) Structural insight into poly(A) binding and catalytic mechanism of human PARN. EMBO J, 24, 4082–4093.

107. Yang, X., Morita, M., Wang, H., Suzuki, T., Yang, W., Luo, Y., Zhao, C., Yu, Y., Bartlam, M., Yamamoto, T., et al. (2008) Crystal structures of human BTG2 and mouse TIS21 involved in suppression of CAF1 deadenylase activity. Nucleic Acids Research, 36, 6872–6881.

108. Miyasaka, T., Morita, M., Ito, K., Suzuki, T., Fukuda, H., Takeda, S., Inoue, J.-I., Semba, K. and Yamamoto, T. (2008) Interaction of antiproliferative protein Tob with the CCR4-NOT deadenylase complex. Cancer Sci, 99, 755–761.

109. Rouault, J.P., Rimokh, R., Tessa, C., Paranhos, G., Ffrench, M., Duret, L., Garoccio, M., Germain, D., Samarut, J. and Magaud, J.P. (1992) BTG1, a member of a new family of antiproliferative genes. EMBO J, 11, 1663–1670.

110. Matsuda, S., Kawamura-Tsuzuku, J., Ohsugi, M., Yoshida, M., Emi, M., Nakamura, Y., Onda, M., Yoshida, Y., Nishiyama, A. and Yamamoto, T. (1996) Tob, a novel protein that interacts with p185erbB2, is associated with anti-proliferative activity. Oncogene, 12, 705–713.

111. Montagnoli, A., Guardavaccaro, D., Starace, G. and Tirone, F. (1996) Overexpression of the nerve growth factor-inducible PC3 immediate early gene is associated with growth inhibition. Cell Growth Differ, 7, 1327–1336.

112. Buanne, P., Corrente, G., Micheli, L., Palena, A., Lavia, P., Spadafora, C., Lakshmana, M.K., Rinaldi, A., Banfi, S., Quarto, M., et al. (2000) Cloning of PC3B, a novel member of the PC3/BTG/TOB family of growth inhibitory genes, highly expressed in the olfactory epithelium. Genomics, 68, 253–263.

113. Doidge, R., Mittal, S., Aslam, A. and Winkler, G.S. (2012) The Anti-Proliferative Activity of BTG/TOB Proteins Is Mediated via the Caf1a (CNOT7) and Caf1b (CNOT8) Deadenylase Subunits of the Ccr4-Not Complex. PLoS ONE, 7.

114. Shi, X., Halder, P., Yavuz, H., Jahn, R. and Shuman, H.A. (2016) Direct targeting of membrane fusion by SNARE mimicry: Convergent evolution of *Legionella* effectors. Proceedings of the National Academy of Sciences of the United States of America, 113, 8807–8812.

115. Price, C.T., Al-Khodor, S., Al-Quadan, T., Santic, M., Habyarimana, F., Kalia, A. and Kwaik, Y.A. (2009) Molecular Mimicry by an F-Box Effector of *Legionella pneumophila* Hijacks a Conserved Polyubiquitination Machinery within Macrophages and Protozoa. PLoS Pathog, 5, e1000704.

116. Coller, J. and Wickens, M. (2002) Tethered function assays using 3’ untranslated regions. Methods, 26, 142–150.

117. Finoux, A.-L. and Séraphin, B. (2006) *In vivo* targeting of the yeast Pop2 deadenylase subunit to reporter transcripts induces their rapid degradation and generates new decay intermediates. Journal of Biological Chemistry, 281, 25940–25947.

118. Luo, E.-C., Nathanson, J.L., Tan, F.E., Schwartz, J.L., Schmok, J.C., Shankar, A., Markmiller, S., Yee, B.A., Sathe, S., Pratt, G.A., et al. (2020) Large-scale tethered function assays identify factors that regulate mRNA stability and translation. Nature Structural & Molecular Biology, 27, 989– 1000.

119. Zipprich, J.T., Bhattacharyya, S., Mathys, H. and Filipowicz, W. (2009) Importance of the C-terminal domain of the human GW182 protein TNRC6C for translational repression. RNA, 15, 781–793.

120. Chapat, C., Chettab, K., Simonet, P., Wang, P., La Grange, De, P., Le Romancer, M. and Corbo, L. (2017) Alternative splicing of CNOT7 diversifies CCR4–NOT functions. Nucleic Acids Research, 45, 8508–8523.

121. Zekri, L., Kuzuoğlu-Öztürk, D. and Izaurralde, E. (2013) GW182 proteins cause PABP dissociation from silenced miRNA targets in the absence of deadenylation. The EMBO Journal, 32, 1052– 1065.

122. Cooke, A., Prigge, A. and Wickens, M. (2010) Translational repression by deadenylases. J. Biol. Chem., 285, 28506–28513.

123. Chekulaeva, M., Mathys, H., Zipprich, J.T., Attig, J., Colic, M., Parker, R. and Filipowicz, W. (2011) miRNA repression involves GW182-mediated recruitment of CCR4-NOT through conserved W-containing motifs. Nature Structural & Molecular Biology, 18, 1218–1226.

124. Bawankar, P., Loh, B., Wohlbold, L., Schmidt, S. and Izaurralde, E. (2013) NOT10 and C2orf29/NOT11 form a conserved module of the CCR4-NOT complex that docks onto the NOT1 N-terminal domain. RNA Biology, 10, 228–244.

125. Aslam, A., Mittal, S., Koch, F., Andrau, J.-C. and Winkler, G.S. (2009) The CCR4-NOT deadenylase subunits CNOT7 and CNOT8 have overlapping roles and modulate cell proliferation. Molecular Biology of the Cell, 20, 3840–3850.

126. Morel, A.P., Sentis, S., Bianchin, C., Le Romancer, M., Jonard, L., Rostam, M.C., Rimokh, R. and Corbo, L. (2003) BTG2 antiproliferative protein interacts with the human CCR4 complex existing *in vivo* in three cell-cycle-regulated forms. Journal of Cell Science, 116, 2929–2936.

127. Maekawa, M., Yamamoto, T. and Nishida, E. (2004) Regulation of subcellular localization of the antiproliferative protein Tob by its nuclear export signal and bipartite nuclear localization signal sequences. Exp Cell Res, 295, 59–65.

128. Denis, C.L., Chiang, Y.C., Cui, Y.J. and Chen, J.J. (2001) Genetic evidence supports a role for the yeast CCR4-NOT complex in transcriptional elongation. Genetics, 158, 627–634.

129. Kruk, J.A., Dutta, A., Fu, J., Gilmour, D.S. and Reese, J.C. (2011) The multifunctional Ccr4–Not complex directly promotes transcription elongation. Genes Dev., 25, 581–593.

130. Reese, J.C. (2013) The control of elongation by the yeast Ccr4-not complex. Biochim. Biophys. Acta, 1829, 127–133.

131. Dutta, A., Babbarwal, V., Fu, J., Brunke-Reese, D., Libert, D.M., Willis, J. and Reese, J.C. (2015) Ccr4-Not and TFIIS Function Cooperatively To Rescue Arrested RNA Polymerase II. Mol. Cell. Biol., 35, 1915–1925.

132. Exinger, F. and Lacroute, F. (1992) 6-Azauracil inhibition of GTP biosynthesis in *Saccharomyces cerevisiae*. Curr. Genet., 22, 9–11.

133. López-Rosas, I., Orozco, E., Marchat, L.A., García-Rivera, G., Guillen, N., Weber, C., Carrillo- Tapia, E., Hernández de la Cruz, O., Pérez-Plasencia, C. and López-Camarillo, C. (2012) mRNA decay proteins are targeted to poly(A)+ RNA and dsRNA-containing cytoplasmic foci that resemble P-bodies in *Entamoeba histolytica*. PLoS ONE, 7, e45966.

134. López-Rosas, I., Marchat, L.A., Olvera, B.G., Guillen, N., Weber, C., Hernández de la Cruz, O., Ruíz-García, E., Astudillo-de la Vega, H. and López-Camarillo, C. (2014) Proteomic analysis identifies endoribouclease EhL-PSP and EhRRP41 exosome protein as novel interactors of EhCAF1 deadenylase. J Proteomics, 111, 59–73.

135. Schwede, A., Ellis, L., Luther, J., Carrington, M., Stoecklin, G. and Clayton, C. (2008) A role for Caf1 in mRNA deadenylation and decay in trypanosomes and human cells. Nucleic Acids Research, 36, 3374–3388.

136. Schwede, A., Manful, T., Jha, B.A., Helbig, C., Bercovich, N., Stewart, M. and Clayton, C. (2009) The role of deadenylation in the degradation of unstable mRNAs in trypanosomes. Nucleic Acids Research, 37, 5511–5528.

137. Erben, E., Chakraborty, C. and Clayton, C. (2014) The CAF1-NOT complex of trypanosomes. Frontiers in Genetics, 4, 299.

138. Sarowar, S., Oh, H.W., Cho, H.S., Baek, K.-H., Seong, E.S., Joung, Y.H., Choi, G.J., Lee, S. and Choi, D. (2007) *Capsicum annuum* CCR4-associated factor CaCAF1 is necessary for plant development and defence response. Plant J, 51, 792–802.

139. Liang, W., Li, C., Liu, F., Jiang, H., Li, S., Sun, J., Wu, X. and Li, C. (2009) The *Arabidopsis* homologs of CCR4-associated factor 1 show mRNA deadenylation activity and play a role in plant defence responses. Cell Research, 19, 307–316.

140. Cernadas, R.A., Camillo, L.R. and Benedetti, C.E. (2008) Transcriptional analysis of the sweet orange interaction with the citrus canker pathogens *Xanthomonas axonopodis* pv. citri and *Xanthomonas axonopodis* pv. aurantifolii. Mol Plant Pathol, 9, 609–631.

141. Hart, K.J., Oberstaller, J., Walker, M.P., Minns, A.M., Kennedy, M.F., Padykula, I., Adams, J.H. and Lindner, S.E. (2019) *Plasmodium* male gametocyte development and transmission are critically regulated by the two putative deadenylases of the CAF1/CCR4/NOT complex. PLoS Pathog, 15, e1007164.

142. Panepinto, J.C., Heinz, E. and Traven, A. (2013) The cellular roles of Ccr4-NOT in model and pathogenic fungi-implications for fungal virulence. Frontiers in Genetics, 4, 302.

143. Jiang, G., Gong, M., Song, H., Sun, W., Zhao, W. and Wang, L. (2020) Tob2 Inhibits TLR-Induced Inflammatory Responses by Association with TRAF6 and MyD88. J. Immunol., 205, 981–986.

144. Hwang, S.S., Lim, J., Yu, Z., Kong, P., Sefik, E., Xu, H., Harman, C.C.D., Kim, L.K., Lee, G.R., Li, H.- B., et al. (2020) mRNA destabilization by BTG1 and BTG2 maintains T cell quiescence. Science (New York, N.Y.), 367, 1255–1260.

145. de Jesús-Díaz, D.A., Murphy, C., Sol, A., Dorer, M. and Isberg, R.R. (2017) Host Cell S Phase Restricts *Legionella pneumophila* Intracellular Replication by Destabilizing the Membrane-Bound Replication Compartment. mBio, 8, 376–18.

146. Harb, O.S., Gao, L.-Y. and Kwaik, Y.A. (2000) From protozoa to mammalian cells: a new paradigm in the life cycle of intracellular bacterial pathogens. Environmental Microbiology, 2, 251–265.

147. Mengue, L., Régnacq, M., Aucher, W., Portier, E., Héchard, Y. and Samba-Louaka, A. (2016) *Legionella pneumophila* prevents proliferation of its natural host *Acanthamoeba castellanii*. Sci. Rep., 6, 36448.

148. Prévôt, D., Morel, A.P., Voeltzel, T., Rostan, M.C., Rimokh, R., Magaud, J.P. and Corbo, L. (2001) Relationships of the antiproliferative proteins BTG1 and BTG2 with CAF1, the human homolog of a component of the yeast CCR4 transcriptional complex: involvement in estrogen receptor alpha signaling pathway. Journal of Biological Chemistry, 276, 9640–9648.

149. Bogdan, J., Adams-Burton, C., Pedicord, D., Sukovich, D., Benfield, P., Corjay, M., Stoltenberg, J. and Dicker, I. (1998) Human carbon catabolite repressor protein (CCR4)-associative factor 1: cloning, expression and characterization of its interaction with the B-cell translocation protein BTG1. Biochemical Journal, 336, 471–481.

150. Chang, H., Lim, J., Ha, M. and Kim, V.N. (2014) TAIL-seq: genome-wide determination of poly(A) tail length and 3’ end modifications. Molecular Cell, 53, 1044–1052.

