## Supplementary material for "The *L. pneumophila* effector PieF modulates mRNA stability through association with eukaryotic CCR4-NOT": Figure S1

**Supplementary Figure**

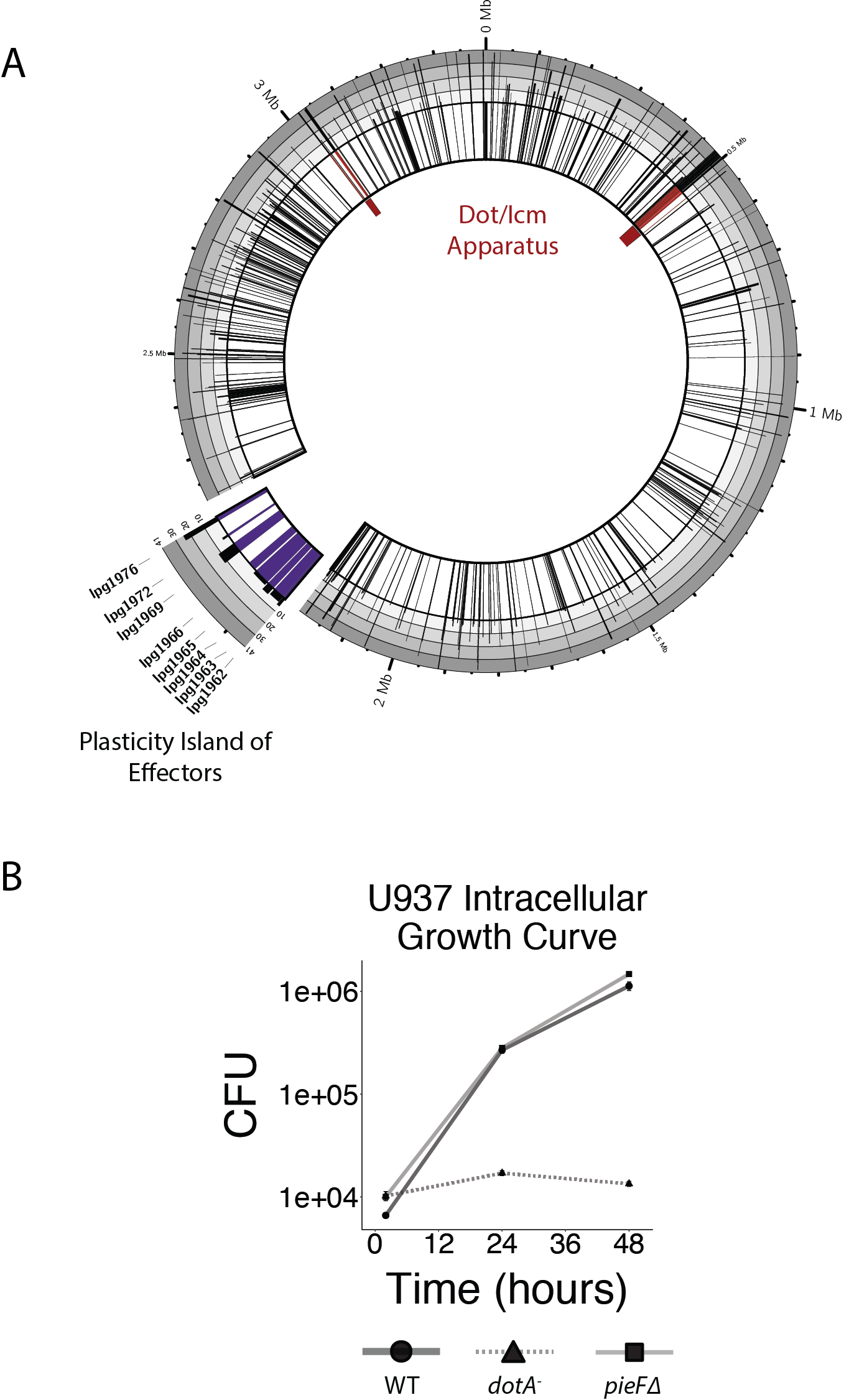

Supplemental Figure 1 A) *PieF* (Lpg1972) is found within the *pie* locus of *L. pneumophila* (Ninio *et al*., 2009). The innermost ring of the Circos plot represents the distribution of effectors across the *L. pneumophila* Str. Philadelphia-1 genome as a barcode. The *pie* locus is emphasized, and *pie* effectors are colored purple. The outermost ring is a bar plot representation of effector conservation across 41 sequenced *Legionella* species (Burstein *et al*., 2016). B) Intracellular replication was measured by CFU plating at three time points (2, 24, and 48 h) during U937 infection (initial MOI 0.05) onto charcoal yeast. Deletion of *pieF* shows no replication defect relative to wildtype, the *dotA*- mutant lacking a functional type IV secretion system fails to replicate intracellularly.
