## Supplementary material for "The *L. pneumophila* effector PieF modulates mRNA stability through association with eukaryotic CCR4-NOT": Table S1

**Supplementary Tables**

Supp. Table 1 Plasmids used in this study.

| Name | Description | Use | Source |
| --- | --- | --- | --- |
| pcDNA3.1-HA-mCherry-ccdB | DB3.1 + pcDNA3.1-HA-mCherry-ccdB | AP-MS, fluorescence microscopy | Gift from Isberg Lab, modified pcDNA-DEST53 |
| pcDNA3.1-HA-mCherry-ccdB | Top10 + HA-mCherry-Lpg1972 | AP-MS, fluorescence microscopy | This study |
| pcDNA3.1-HA-mCherry-ccdB | Top10 + HA-mCherry-Lpg2149 | AP-MS, fluorescence microscopy | This study |
| pKT25 | Top10 + pKT25 | B2H | Karimova et al. 1998, kind gift from Dr. K. Maxwell |
| pUT18C | Top10 + pUT18C | B2H | Karimova et al. 1998, kind gift from Dr. K. Maxwell |
| pKT25-Lpg1972 | Top10 + pKT25-Lpg1972 | B2H | This study |
| pUT18C-CNOT2 | Top10 + pUT18C-CNOT2 | B2H | This study |
| pUT18C-CNOT3 | Top10 + pUT18C-CNOT3 | B2H | This study |
| pUT18C-CNOT4 | Top10 + pUT18C-CNOT4 | B2H | This study |
| pUT18C-CNOT6 | Top10 + pUT18C-CNOT6 | B2H | This study |
| pUT18C-CNOT6L | Top10 + pUT18C-CNOT6L | B2H | This study |
| pUT18C-CNOT7 | Top10 + pUT18C-CNOT7 | B2H | This study |
| pUT18C-CNOT8 | Top10 + pUT18C-CNOT8 | B2H | This study |
| pUT18C-CNOT9 | Top10 + pUT18C-CNOT9 | B2H | This study |
| pUT18C-CNOT10 | Top10 + pUT18C-CNOT10 | B2H | This study |
| pUT18C-CNOT11 | Top10 + pUT18C-CNOT11 | B2H | This study |
| pUT18C-CNOT1 | Top10 + pUT18C-CNOT1 | B2H | This study |
| pMSGC68-SBP-TEV | Top10 + pMSGC68-SBP-TEV | Cloning | DNASU ID:528, Gift from A. Savchenko |
| pJB4648-Lpg1972 | Top10 + Lpg1972-deletion construct L.P. | Deletion in *L. pneumophila* | This study |
| R600K | R600K | Deletion in *L. pneumophila* | Swanson & Isberg 1996 |
| pT7-EGFP-C1-HsNot7 | Top10 + pT7-EGFP-C1-HsNot7 | Fluorescence microscopy | pT7-EGFP-C1-HsNot7 was a gift from Elisa Izaurralde (Addgene plasmid # 37325 ; http://n2t.net/addgene:37325 ; RRID:Addgene_37325 |
| pMSGC68 | BL21 + pMSGC68-CNOT7 | Protein Purification | This study |
| pOG44 | Top10 + pOG44 | Strain construction | ThermoFisher V600520 |
| CNOT7-3X-FLAG | Top10 + CNOT7-3X-FLAG Stable Cell Integration | Strain construction | This study |
| FL | Top10 + Firefly luciferase reporter | Tethering | Chapat et al. 2017, Kind gift from Dr. M. Fabian |
| RL-5BoxB | Top10 + Renilla Luciferase with 5X-BoxB 3' UTR | Tethering | Chapat et al. 2017, Kind gift from Dr. M. Fabian |
| RL-5BoxB-A114-N40-HhR | Top10 + Firefly Luciferase with 5X-BoxB hammerhead poly(A)40 3' UTR | Tethering | Chapat et al. 2017, Kind gift from Dr. M. Fabian |
| pCI-lambda-NHA | Top10 + lambda-N-HA-LacZ | Tethering | Nishimura et al. 2018, Kind gift from Dr. M. Fabian |
| pCI-lambda-NHA | Top10 + lambda-N-HA-Lpg1972 | Tethering | This study |
| pCI-lambda-NHA | Top10 + lambda-N-HA-CNOT7 | Tethering | This study |
| pCI-lambda-NHA | Top10 + Lambda-N-HA-BTG1 | Tethering | This study |
| pDEST_AD_Song_V3-EV | DB3.1 + AD-Y2H-EV | Y2H | Yachie et al. 2016 |
| pDEST_DB_Song_V3-EV | DB3.1 + DB-Y2H-EV | Y2H | Yachie et al. 2016 |
| pDEST_DB_Song_V3-Lpg1972 | Top10 + DB-Y2H-Lpg1972 | Y2H | This study |
| pDEST_AD_Song_V3-CNOT7 | Top10 + AD-Y2H-CNOT7 | Y2H | This study |
| pDEST_AD_Song_V3-CNOT1 | Top10 + AD-Y2H-CNOT1 | Y2H | This study |
| pDEST_AD_Song_V3-CNOT2 | Top10 + AD-Y2H-CNOT2 | Y2H | This study |
| pDEST_AD_Song_V3-CNOT6 | Top10 + AD-Y2H-CNOT6 | Y2H | This study |
| pDEST_AD_Song_V3-CNOT6L | Top10 + AD-Y2H-CNOT6L | Y2H | This study |
| pDEST_AD_Song_V3-CNOT8 | Top10 + AD-Y2H-CNOT8 | Y2H | This study |
| pDEST_AD_Song_V3-CNOT9 | Top10 + AD-Y2H-CNOT9 | Y2H | This study |
| pDEST_AD_Song_V3-CNOT10 | Top10 + AD-Y2H-CNOT10 | Y2H | This study |
| pDEST_AD_Song_V3-CNOT3 | Top10 + AD-Y2H-CNOT3 | Y2H | This study |
| pDEST_DB_Song_V3-CNOT6L | Top10 + DB-Y2H-CNOT6L | Y2H | This study |
| pDEST_AD_Song_V3-CNOT4 | Top10 + AD-Y2H-CNOT4 | Y2H | This study |
| pDEST_AD_Song_V3-CNOT11 | Top10 + AD-Y2H-CNOT11 | Y2H | This study |
| pDEST_AD_Song_V3-CNOT7[Y260A, E247A] | Top10 + AD-Y2H-CNOT7[Y260A, E247A] | Y2H | This study |
| pDEST_DB_Song_V3-BTG1 | Top10 + DB-Y2H-BTG1 | Y2H | This study |
| pAG416-GpdP-Lpg1972 | Top10 + pAG416-GpdP-Lpg1972 | Yeast growth curves | This study |
| pAG416-GpdP-Lpg2149 | Top10 + pAG416-GpdP-Lpg2149 | Yeast growth curves | This study |
| pAG416-GpdP-CcdB | DB3.1 + pAG416-GpdP-EV | Yeast growth curves | Alberti et al. 2007 |
