## Supplementary material for "The *L. pneumophila* effector PieF modulates mRNA stability through association with eukaryotic CCR4-NOT": Table S2

**Supplementary Tables**

Table S2 Bacterial Strains used in this study.

| Name | Genotype | Use | Source |
| --- | --- | --- | --- |
| BTH101 | F- , cya-99, araD139, galE15, galK16, rpsL1 (Str r ), hsdR2, mcrA1, mcrB1. | B2H | Karimova et al.1998, Kind gift from Dr. K. Maxwell |
| Top10 | F–*mcr*A Δ(*mrr*-*hsd*RMS-*mcr*BC) φ80*lac*ZΔM15 Δ*lac*X74 *rec*A1 *ara*D139 Δ(*ara-leu*)7697 *gal*U *gal*K λ–*rps*L(StrR) *end*A1 *nup*G | Cloning | ThermoFisher Scientific C404010 |
| BL21 (DE3) | F–ompT hsdSB (rB–, mB–) gal dcm (DE3) | Protein Purification | ThermoFisher Scientific EC0114 |
| DB3.1 | F–*mcr*A Δ(*mrr*-*hsd*RMS-*mcr*BC) φ80*lac*ZΔM15 Δ*lac*X74 *rec*A1 *ara*Δ139 Δ(*ara*-*leu*)7697 *gal*U *gal*K *rps*L(StrR) *end*A1 *nup*G *fhu*A::*IS2* | Cloning | ThermoFisher Scientific E7650 |
| XL1Blue | : recA1 endA1 gyrA96 thi-1 hsdR17 supE44 relA1 lac [F´ proAB lacIq Z∆M15 Tn10 (Tetr )] | Cloning | Agilent 200249 |
| BL21 + pMGCS68-Lpg1972 | see BL21(DE3) | Protein Purification | This study |
| XL1Blue + pKT25 (bacterial two-hybrid EV) | see XL1Blue | Cloning | Karimova et al. 1998 |
| XL1Blue + pUT18C (bacterial two-hybrid EV) | see XL1Blue | Cloning | Karimova et al. 1998 |
| BTH101 + pKT25-EV + pUT18C-EV | see BTH101 | B2H | This study |
| BTH101 + pKT25-EV + pUT18C-CNOT6 | see BTH101 | B2H | This study |
| BTH101 + pKT25-EV + pUT18C-CNOT7 | see BTH101 | B2H | This study |
| BTH101 + pKT25-Lpg1972 + pUT18C-EV | see BTH101 | B2H | This study |
| BTH101 + pKT25-Lpg1972 + pUT18C-CNOT6 | see BTH101 | B2H | This study |
| BTH101 + pKT25-Lpg1972 + pUT18C-CNOT7 | see BTH101 | B2H | This study |
| BTH101 + pKT25-EV + pUT18C-CNOT2 | see BTH101 | B2H | This study |
| BTH101 + pKT25-EV + pUT18C-CNOT3 | see BTH101 | B2H | This study |
| BTH101 + pKT25-EV + pUT18C-CNOT4 | see BTH101 | B2H | This study |
| BTH101 + pKT25-EV + pUT18C-CNOT6L | see BTH101 | B2H | This study |
| BTH101 + pKT25-EV + pUT18C-CNOT8 | see BTH101 | B2H | This study |
| BTH101 + pKT25-EV + pUT18C-CNOT9 | see BTH101 | B2H | This study |
| BTH101 + pKT25-EV + pUT18C-CNOT10 | see BTH101 | B2H | This study |
| BTH101 + pKT25-EV + pUT18C-CNOT11 | see BTH101 | B2H | This study |
| BTH101 + pKT25-Lpg1972 + pUT18C-CNOT1 | see BTH101 | B2H | This study |
| BTH101 + pKT25-Lpg1972 + pUT18C-CNOT2 | see BTH101 | B2H | This study |
| BTH101 + pKT25-Lpg1972 + pUT18C-CNOT3 | see BTH101 | B2H | This study |
| BTH101 + pKT25-Lpg1972 + pUT18C-CNOT4 | see BTH101 | B2H | This study |
| BTH101 + pKT25-Lpg1972 + pUT18C-CNOT6L | see BTH101 | B2H | This study |
| BTH101 + pKT25-Lpg1972 + pUT18C-CNOT8 | see BTH101 | B2H | This study |
| BTH101 + pKT25-Lpg1972 + pUT18C-CNOT9 | see BTH101 | B2H | This study |
| BTH101 + pKT25-Lpg1972 + pUT18C-CNOT10 | see BTH101 | B2H | This study |
| BTH101 + pKT25-Lpg1972 + pUT18C-CNOT11 | see BTH101 | B2H | This study |
| BTH101 + pKT25 EV + pUT18C CNOT1 | see BTH101 | B2H | This study |
| BL21 + pMSGC68-CNOT7 | see BL21(DE3) | Protein Purification | This study |
