## Supplementary material for "The *L. pneumophila* effector PieF modulates mRNA stability through association with eukaryotic CCR4-NOT": Table S3

**Supplementary Tables**

Supp. Table 3 *L. pneumophila* strains used in this study.

| Accession | Alias | Source |
| --- | --- | --- |
| **hLP01** | LP02 | Berger *et al.* 1993 |
| **hLP02** | LP03 | Berger *et al.* 1993 |
| **hLP03** | LP02 1972Δ | this study |
