## Supplementary material for "The *L. pneumophila* effector PieF modulates mRNA stability through association with eukaryotic CCR4-NOT": Table S4

**Supplementary Tables**

Supp. Table 4. Primers & Oligonucleotide(s) used in this study

| Accession | Name | Sequence (5′-3′) | Use | Source |
| --- | --- | --- | --- | --- |
| oHM389 | pKT25_fwd_screen | tatgccgcatctgtccaacttcc | B2H | This study |
| oHM390 | pUT18C_fwd_screen | ttctcgccggatgtactggaaacg | B2H | This study |
| oHM408 | 1972_fwd_pKT25 | ggcgggctgcagggtcgactatgaaaagactaattatctgtaatg | B2H | This study |
| oHM409 | 1972_rev_pKT25 | cttaggtacccggggatccttcaaattttgtttattaagttatatgc | B2H | This study |
| oHM438 | CNOT1_fwd_pUT18C | aacgccactgcaggtcgactatgaatcttgactcgctctc | B2H | This study |
| oHM439 | CNOT1_rev_pUT18C | gctcggtacccggggatcctctaactggcacctgtccc | B2H | This study |
| oHM440 | CNOT2_fwd_pUT18C | aacgccactgcaggtcgactatggtgaggactgatggac | B2H | This study |
| oHM441 | CNOT2_rev_pUT18C | gctcggtacccggggatcctttagaaggcttgctgagc | B2H | This study |
| oHM442 | CNOT3_fwd_pUT18C | aacgccactgcaggtcgactatggcggacaagcgcaaac | B2H | This study |
| oHM443 | CNOT3_rev_pUT18C | gctcggtacccggggatccttcactggaggtcccggtc | B2H | This study |
| oHM444 | CNOT4_fwd_pUT18C | aacgccactgcaggtcgactatgtctcgcagtcctgatg | B2H | This study |
| oHM445 | CNOT4_rev_pUT18C | gctcggtacccggggatccttcaggccacagtagtgtg | B2H | This study |
| oHM446 | CNOT6_fwd_pUT18C | aacgccactgcaggtcgactatgcccaaagaaaaatacgag | B2H | This study |
| oHM447 | CNOT6_rev_pUT18C | gctcggtacccggggatcctctacctcctgccaggaag | B2H | This study |
| oHM448 | CNOT6L_fwd_pUT18C | aacgccactgcaggtcgactatgagactaatagggatgc | B2H | This study |
| oHM449 | CNOT6L_rev_pUT18C | gctcggtacccggggatcctctacctccgattaggcaag | B2H | This study |
| oHM450 | CNOT7_fwd_pUT18C | aacgccactgcaggtcgactatgccagcggcaactgtag | B2H | This study |
| oHM451 | CNOT7_rev_pUT18C | gctcggtacccggggatccttcatgactgcttgttggcttc | B2H | This study |
| oHM452 | CNOT8_fwd_pUT18C | aacgccactgcaggtcgactatgcctgcagcacttgtg | B2H | This study |
| oHM453 | CNOT8_rev_pUT18C | gctcggtacccggggatccttcactgctgcatgttgttg | B2H | This study |
| oHM454 | CNOT9_fwd_pUT18C | aacgccactgcaggtcgactatgcacagcctggcgacg | B2H | This study |
| oHM455 | CNOT9_rev_pUT18C | gctcggtacccggggatccttcactgagggggcagggg | B2H | This study |
| oHM456 | CNOT10_fwd_pUT18C | aacgccactgcaggtcgactatggctgcagacaagcctg | B2H | This study |
| oHM457 | CNOT10_rev_pUT18C | gctcggtacccggggatccttcactttctctgcacagtgg | B2H | This study |
| oHM458 | CNOT11_fwd_pUT18C | aacgccactgcaggtcgactatgcccggcggaggggcg | B2H | This study |
| oHM459 | CNOT11_rev_pUT18C | gctcggtacccggggatcctttattttgacatttcggtctcagaaggtgtttccccagtatccaatgtcttc | B2H | This study |
| oHM391 | CNOT1_rev_internal | gcgtctatctgttcttgtccaactcc | B2H, tethering | This study |
| oHM392 | CNOT2_rev_internal | aactgctcatgctatttgtcctgc | B2H, tethering | This study |
| oHM393 | CNOT3_rev_internal | gcctcttgtccttgatctcgttgg | B2H, tethering | This study |
| oHM394 | CNOT4_rev_internal | atcggcaaatctggtagccacagg | B2H, tethering | This study |
| oHM395 | CNOT6_rev_internal | tttccgagttctgcgggtaagc | B2H, tethering | This study |
| oHM396 | CNOT6L_rev_internal | tttcccattggctacctcctctgc | B2H, tethering | This study |
| oHM397 | CNOT7_rev_internal | tgcaaccacacctggaaactcg | B2H, tethering | This study |
| oHM398 | CNOT8_rev_internal | ctggcccacacttcacagataacc | B2H, tethering | This study |
| oHM400 | CNOT10_rev_internal | catgtttctctgctccctgatctgc | B2H, tethering | This study |
| oHM401 | CNOT11_rev_internal | aagtggtcggccttgctgaagtag | B2H, tethering | This study |
| oHM402 | Lpg1972_rev_internal | ttgcctgagaagaggttagctcc | B2H, tethering | This study |
| oHM402 | CNOT9_rev_internal | ttctcgcttcttacttagctcc | B2H, tethering | This study |
| oHM89 | M13 fwd | gtaaaacgacggccagt | Cloning | ThermoFisher Scientific N52002 |
| oHM90 | M13 rev | ggaaacagctatgaccatg | Cloning | ThermoFisher Scientific N53002 |
| oHM312 | CNOT7_attB_F | ggggacaagtttgtacaaaaaagcaggcttcatgccagcggcaactgtagatcata | Cloning | This study |
| oHM313 | CNOT7_attB_R | ggggaccactttgtacaagaaagctgggtcctatcatgactgcttgttggcttcctct | Cloning | This study |
| oHM351 | Lpg1972_attB_Fwd | ggggacaagtttgtacaaaaaagcaggcttcatgaaaagactaattatctgtaatg | Cloning | This study |
| oHM352 | Lpg1972_attB_REV_NO_STOP | ggggaccactttgtacaagaaagctgggtcctaaattttgtttattaagttatatgct | Cloning | This study |
| oHM509 | BTG1_attB_fwd | ggggacaagtttgtacaaaaaagcaggcttcatgcatcccttctacacccgggccg | Cloning | This study |
| oHM510 | BTG1_attB_rev | ggggaccactttgtacaagaaagctgggtcctattaacctgatacagtcatcatattg | Cloning | This study |
| oHM480 | 6-FAM-RNA-poly(A)-20 | ucuaaauaaaaaaaaaaaaaaaaaaaa | Deadenylation assays | Wang et al. 2010 |
| oHM263 | Lpg1972-SacI-UpF_-379 | gagctcttgaaagcatgatcccggtc | Gene deletion | This study |
| oHM264 | Lpg1972-SeqF | tgcaggccaaacactgagtc | Gene deletion | This study |
| oHM265 | Lpg1972-dsF | tctaaaatgtgagctaactataagctatccaggccaagga | Gene deletion | This study |
| oHM266 | Lpg1972-upR | tccttggcctggatagcttatagttagctcacattttaga | Gene deletion | This study |
| oHM267 | Lpg1972-SeqR | cactttaattcgcgcctgg | Gene deletion | This study |
| oHM268 | Lpg1972-SalI-dsR | gtcgaccctcggaagtcatggaaacatc | Gene deletion | This study |
| oHM476 | Lpg1972_LIC_Fwd_PMSG68 | tacttccaatccaatgccatgaaaagactaattatct | Protein Purification | This study |
| oHM477 | Lpg1972_LIC_Rev_PMSG68 | ttatccacttccaatgttatcaaattttgtttattaagt | Protein Purification | This study |
| oHM478 | CNOT7_LIC_Fwd_PMSG68 | tacttccaatccaatgccatgccagcggcaactgtag | Protein Purification | This study |
| oHM479 | CNOT7_LIC_Rev_PMSG68 | ttatccacttccaatgttatgactgcttgttggcttcc | Protein Purification | This study |
| oHM489 | 1972_tethering_amp_F | actcgacgcggatcccgtcgaattcatgaaaagactaattatctgtaatg | Tethering | This study |
| oHM490 | 1972_tethering_amp_R | aaccctcactaaagggaagcggccgctcaaattttgtttattaagttatatgc | Tethering | This study |
| oHM491 | CNOT7_tethering_amp_F | actcgacgcggatcccgtcgaattcatgccagcggcaactgtag | Tethering | This study |
| oHM492 | CNOT7_tethering_amp_R | aaccctcactaaagggaagcggccgctcatgactgcttgttggcttc | Tethering | This study |
| oHM495 | LacZ_tethering_amp_F | actcgacgcggatcccgtcgaattcatgaccatgattacggattc | Tethering | This study |
| oHM496 | LacZ_tethering_amp_R | aaccctcactaaagggaagcggccgcttatttttgacaccagacc | Tethering | This study |
| oHM497 | RL_fwd_qPCR | aatggctcatatcgcctcctgg | Tethering | This study |
| oHM498 | RL_rev_qPCR | cttgatcttgtcttggtgctcgtagg | Tethering | This study |
| oHM499 | FL_fwd_qPCR | aggctatgaagagatacgccctgg | Tethering | This study |
| oHM500 | FL_rev_qPCR | ctgcaactccgataaataacgcgc | Tethering | This study |
| oHM503 | pCLNeo-LambdaN-HA-Seq-Fwd | tttctctccacaggtgtccactcc | Tethering | This study |
| oHM504 | pCLNeo-LambdaN-HA-Seq-Rev | ggaagggaagaaagcgaaaggagc | Tethering | This study |
| oHM506 | LacZ-internal-rev | gataggttacgttggtgtagatgggc | Tethering | This study |
| oHM511 | BTG1_pCLNeo_F | actcgacgcggatcccgtcgaattcatgcatcccttctacacccgggccg | Tethering | This study |
| oHM512 | BTG1_pCLNeo_R | aaccctcactaaagggaagcggccgcttaacctgatacagtcatcatattg | Tethering | This study |
| oHM519 | CNOT7_Y260A_fwd | tggtcatttggccggccttggttc | Y2H | This study |
| oHM520 | CNOT7_Y260A_rev | caatatttggcatcatcaatatg | Y2H | This study |
| oHM521 | CNOT7_E247A_fwd | aatgttctttgccgatcatattgatgatgc | Y2H | This study |
| oHM522 | CNOT7_E247A_rev | tctctcattttgaaaaaggc | Y2H | This study |
