## Supplementary material for "The *L. pneumophila* effector PieF modulates mRNA stability through association with eukaryotic CCR4-NOT": Table S5

**Supplementary Tables**

Supp. Table 5 Yeast strains used in this study.

| Name | Use | Genotype | Source |
| --- | --- | --- | --- |
| RY1010 | iBFG-Y2H | MATa *leu2-3,112 trp1-901 his3-200 ura3-52 gal4∆ gal80∆ PGAL2-ADE2 LYS2::PGAL1-HIS3 MET2::PGAL7-lacZ cyh2R* *can1∆::PCMV-rtTA-KanMX4* | Yachie et al. 2016 |
| RY1030 | iBFG-Y2H | *MATα Leu2-3,112 trp1-901 his3-200 ura3-52 gal4∆ gal80∆ PGAL2-ADE2 LYS2::PGAL1-HIS3 MET2::PGAL7-lacZ cyh2R* *can1∆::TADH1-PtetO2-Cre-TCYC1-KanMX4* | Yachie et al. 2016 |
| RY1010 + pDEST_AD_Song_V3-EV | Y2H | See Ry1010 | This study |
| RY1030 + pDEST_DB_Song_V3-EV | Y2H | See RY1030 | This study |
| RY1010-1030 + pDEST_AD_Song_V3-EV & pDEST_DB_Song_V3-EV | Y2H | See RY1010 & RY1030 | This study |
| RY1030 + pDEST_DB_Song_V3-Lpg1972 | Y2H | See RY1030 | This study |
| RY1010-1030 + pDEST_AD_Song_V3-EV & Lpg1972-DB Y2H | Y2H | See RY1010 & RY1030 | This study |
| RY1010-1030 + pDEST_AD_Song_V3-CNOT7 & pDEST_DB_Song_V3-EV | Y2H | See RY1010 & RY1030 | This study |
| RY1010-1030 + pDEST_AD_Song_V3-CNOT7 & pDEST_DB_Song_V3-Lpg1972 | Y2H | See RY1010 & RY1030 | This study |
| RY1010 + pDEST_AD_Song_V3-CNOT1 | Y2H | See Ry1010 | This study |
| RY1010 + pDEST_AD_Song_V3-CNOT2 | Y2H | See Ry1010 | This study |
| RY1010 + pDEST_AD_Song_V3-CNOT6 | Y2H | See Ry1010 | This study |
| RY1010 + pDEST_AD_Song_V3-CNOT6L | Y2H | See Ry1010 | This study |
| RY1010 + pDEST_AD_Song_V3-CNOT8 | Y2H | See Ry1010 | This study |
| RY1010 + pDEST_AD_Song_V3-CNOT9 | Y2H | See Ry1010 | This study |
| RY1010 + pDEST_AD_Song_V3-CNOT10 | Y2H | See Ry1010 | This study |
| RY1010 + pDEST_AD_Song_V3-CNOT3 | Y2H | See Ry1010 | This study |
| RY1010-1030 + pDEST_AD_Song_V3-CNOT1 & pDEST_DB_Song_V3-EV | Y2H | See RY1010 & RY1030 | This study |
| RY1010-1030 + pDEST_AD_Song_V3-CNOT1 & pDEST_DB_Song_V3-Lpg1972 | Y2H | See RY1010 & RY1030 | This study |
| RY1010-1030 + pDEST_AD_Song_V3-CNOT2 & pDEST_DB_Song_V3-EV | Y2H | See RY1010 & RY1030 | This study |
| RY1010-1030 + pDEST_AD_Song_V3-CNOT6 & pDEST_DB_Song_V3-EV | Y2H | See RY1010 & RY1030 | This study |
| RY1010-1030 + pDEST_AD_Song_V3-CNOT6L & pDEST_DB_Song_V3-EV | Y2H | See RY1010 & RY1030 | This study |
| RY1010-1030 + pDEST_AD_Song_V3-CNOT8 & pDEST_DB_Song_V3-EV | Y2H | See RY1010 & RY1030 | This study |
| RY1010-1030 + pDEST_AD_Song_V3-CNOT9 & pDEST_DB_Song_V3-EV | Y2H | See RY1010 & RY1030 | This study |
| RY1010-1030 + pDEST_AD_Song_V3-CNOT10 & pDEST_DB_Song_V3-EV | Y2H | See RY1010 & RY1030 | This study |
| RY1010-1030 + pDEST_AD_Song_V3-CNOT3 & pDEST_DB_Song_V3-EV | Y2H | See RY1010 & RY1030 | This study |
| RY1010-1030 + pDEST_AD_Song_V3-CNOT2 & pDEST_DB_Song_V3-Lpg1972 | Y2H | See RY1010 & RY1030 | This study |
| RY1010-1030 + pDEST_AD_Song_V3-CNOT6 & pDEST_DB_Song_V3-Lpg1972 | Y2H | See RY1010 & RY1030 | This study |
| RY1010-1030 + pDEST_AD_Song_V3-CNOT6L & pDEST_DB_Song_V3-Lpg1972 | Y2H | See RY1010 & RY1030 | This study |
| RY1010-1030 + pDEST_AD_Song_V3-CNOT8 & pDEST_DB_Song_V3-Lpg1972 | Y2H | See RY1010 & RY1030 | This study |
| RY1010-1030 + pDEST_AD_Song_V3-CNOT9 & pDEST_DB_Song_V3-Lpg1972 | Y2H | See RY1010 & RY1030 | This study |
| RY1010-1030 + pDEST_AD_Song_V3-CNOT10 & pDEST_DB_Song_V3-Lpg1972 | Y2H | See RY1010 & RY1030 | This study |
| RY1010-1030 + pDEST_AD_Song_V3-CNOT3 & pDEST_DB_Song_V3-Lpg1972 | Y2H | See RY1010 & RY1030 | This study |
| RY1030 + pDEST_DB_Song_V3-CNOT6L | Y2H | See RY1030 | This study |
| RY1010-1030 + pDEST_AD_Song_V3-EV & pDEST_DB_Song_V3-CNOT6L | Y2H | See RY1010 & RY1030 | This study |
| RY1010-1030 + pDEST_AD_Song_V3-CNOT7 & pDEST_DB_Song_V3-CNOT6L | Y2H | See RY1010 & RY1030 | This study |
| RY1010-1030 + pDEST_AD_Song_V3-CNOT7 & pDEST_DB_Song_V3-CNOT6L & pAG416-GpdP-EV | Y2H | See RY1010 & RY1030 | This study |
| RY1010-1030 + pDEST_AD_Song_V3-CNOT7 & pDEST_DB_Song_V3-CNOT6L & pAG416-GpdP-Lpg1972 | Y2H | See RY1010 & RY1030 | This study |
| RY1010-1030 + pDEST_AD_Song_V3-CNOT7 & pDEST_DB_Song_V3-CNOT6L & pAG416-GpdP-Lpg2149 | Y2H | See RY1010 & RY1030 | This study |
| RY1010 + pDEST_AD_Song_V3-CNOT4 | Y2H | See Ry1010 | This study |
| RY1010 + pDEST_AD_Song_V3-CNOT11 | Y2H | See Ry1010 | This study |
| BY4741 | yeast growth curves | MATa his3Δ1 leu2Δ0 met15Δ0 ura3Δ0 | Winston et al. 1995 |
| BY4741 + pAG416-GpdP-EV | yeast growth curves | see BY4741 | This study |
| BY4741 + pAG416-GpdP-Lpg1972 | yeast growth curves | see BY4741 | This study |
| BY4741 + pAG416-GpdP-Lpg2149 | yeast growth curves | see BY4741 | This study |
| RY1010-1030 + pDEST_AD_Song_V3-CNOT4 & pDEST_DB_Song_V3-EV | Y2H | See RY1010 & RY1030 | This study |
| RY1010-1030 + pDEST_AD_Song_V3-CNOT4 & pDEST_DB_Song_V3-Lpg1972 | Y2H | See RY1010 & RY1030 | This study |
| RY1010-1030 + pDEST_AD_Song_V3-CNOT11 & pDEST_DB_Song_V3-EV | Y2H | See RY1010 & RY1030 | This study |
| RY1010-1030 + pDEST_AD_Song_V3-CNOT11 & pDEST_DB_Song_V3-Lpg1972 | Y2H | See RY1010 & RY1030 | This study |
| BY4741 POP2::KanMX | yeast growth curves | MATa his3Δ1 leu2Δ0 met15Δ0 ura3Δ0 pop2::KanMX | Giaever et al. 2002 |
| BY4741 POP2∷KanMX + pAG416-GpdP-CcdB | yeast growth curves | See BY4741 POP2::KanMX | This study |
| RY1010-1030 + pDEST_AD_Song_V3-CNOT7 pDEST_DB_Song_V3-EV; pAG416-GpdP-EV | Y2H | See RY1010 & RY1030 | This study |
| RY1010-1030 + EV-AD CNOT6L-DB; pAG416-GpdP-EV | Y2H | See RY1010 & RY1030 | This study |
| RY1010-1030 + CNOT7[Y260A, pDEST_AD_Song_V3-E247A] & pDEST_DB_Song_V3-EV | Y2H | See RY1010 & RY1030 | This study |
| RY1010-1030 + CNOT7[Y260A, pDEST_AD_Song_V3-E247A] & pDEST_DB_Song_V3-Lpg1972 | Y2H | See RY1010 & RY1030 | This study |
| RY1010-1030 + pDEST_AD_Song_V3-EV & pDEST_DB_Song_V3-BTG1 | Y2H | See RY1010 & RY1030 | This study |
| RY1010-1030 + pDEST_AD_Song_V3-CNOT7 & pDEST_DB_Song_V3-BTG1 | Y2H | See RY1010 & RY1030 | This study |
| RY1010-1030 + CNOT7[Y260A, pDEST_AD_Song_V3-E247A] & pDEST_DB_Song_V3-BTG1 | Y2H | See RY1010 & RY1030 | This study |
