## Supplementary material for "The *L. pneumophila* effector PieF modulates mRNA stability through association with eukaryotic CCR4-NOT": Table S6

**Supplementary Tables**
Supp Table 6. Mammalian cell lines used in this study

| Accession | Alias | Genotype | Source |
| --- | --- | --- | --- |
| HMHs14 | HEK293 T-REx-Flp-In + FLAG EV | FRT::FLAG EV | This study |
| HMHs18 | HEK293 T-REx-Flp-In + FLAG-CNOT7 | FRT::3'FLAG CNOT7 | This study |
| N/A | HEK293T | WT | Gift from Frappier Lab |
| N/A | U937 | WT | Gift from Savchenko Lab |
| N/A | THP1 | WT | Gift from Gray-Owen Lab |
| N/A | HeLa | WT | Gift from Frappier Lab |
