## Supplementary material for "The *L. pneumophila* effector PieF modulates mRNA stability through association with eukaryotic CCR4-NOT": Table S7

**Supplementary Tables**

Supp Table 7. Proteins removed by ProHits artifact, keratin, and ribosomal biofilters.

| Prey Name | Prey Gene ID | Prey Protein ID | Unique Peptides | Bait Name |
| --- | --- | --- | --- | --- |
| TUBB | 203068 | ENSP00000447729 | 100 | CTRL |
| TUBA1B | 10376 | ENSP00000336799 | 98 | CTRL |
| PRPF8 | 10594 | ENSP00000304350 | 92 | CTRL |
| SF3B1 | 23451 | ENSP00000335321 | 50 | CTRL |
| ATP5A1 | 498 | ENSP00000381736 | 45 | CTRL |
| MYH9 | 4627 | ENSP00000216181 | 45 | CTRL |
| EEF1A1 | 1915 | ENSP00000339063 | 44 | CTRL |
| CLTC | 1213 | ENSP00000269122 | 40 | CTRL |
| DHX9 | 1660 | ENSP00000356520 | 34 | CTRL |
| FLNA | 2316 | ENSP00000353467 | 33 | CTRL |
| KRT1 | 3848 | K2C1_HUMAN | 32 | CTRL |
| HNRNPU | 3192 | ENSP00000283179 | 32 | CTRL |
| HSP90AB1 | 3326 | ENSP00000360709 | 29 | CTRL |
| RPS14 | 6208 | ENSP00000385958 | 28 | CTRL |
| SF3B3 | 23450 | ENSP00000305790 | 27 | CTRL |
| HSPA8 | 3312 | ENSP00000437125 | 26 | CTRL |
| KRT9 | 3857 | ENSP00000246662 | 25 | CTRL |
| ACTB | 60 | ENSP00000349960 | 25 | CTRL |
| HSPD1 | 3329 | ENSP00000373620 | 23 | CTRL |
| KRT10 | 3858 | K1C10_HUMAN | 20 | CTRL |
| TRIM28 | 10155 | ENSP00000253024 | 19 | CTRL |
| ATP5B | 506 | ENSP00000262030 | 19 | CTRL |
| HSPA9 | 3313 | ENSP00000297185 | 19 | CTRL |
| RPS3 | 6188 | ENSP00000434643 | 17 | CTRL |
| RPL5 | 6125 | ENSP00000359345 | 17 | CTRL |
| KRT2 | 3849 | ENSP00000310861 | 17 | CTRL |
| DDX17 | 10521 | ENSP00000380033 | 17 | CTRL |
| HNRNPA1 | 3178 | ENSP00000341826 | 16 | CTRL |
| NCL | 4691 | ENSP00000318195 | 16 | CTRL |
| SYNCRIP | 10492 | ENSP00000347380 | 16 | CTRL |
| RPS5 | 6193 | ENSP00000196551 | 15 | CTRL |
| RPS15 | 6209 | ENSP00000233609 | 15 | CTRL |
| RPL9 | 6133 | ENSP00000346022 | 15 | CTRL |
| RPL7 | 6129 | ENSP00000339795 | 15 | CTRL |
| RUVBL2 | 10856 | ENSP00000221413 | 15 | CTRL |
| RPS4X | 6191 | ENSP00000362744 | 14 | CTRL |
| HNRNPM | 4670 | ENSP00000325376 | 14 | CTRL |
| RPL3 | 6122 | ENSP00000346001 | 13 | CTRL |
| RPL7A | 6130 | ENSP00000361076 | 13 | CTRL |
| RPL6 | 6128 | ENSP00000202773 | 13 | CTRL |
| RPS3A | 6189 | ENSP00000346050 | 13 | CTRL |
| DHX15 | 1665 | ENSP00000336741 | 13 | CTRL |
| SFPQ | 6421 | ENSP00000349748 | 13 | CTRL |
| HSPA5 | 3309 | ENSP00000324173 | 13 | CTRL |
| NONO | 4841 | ENSP00000276079 | 12 | CTRL |
| HNRNPA3 | 220988 | ENSP00000376309 | 12 | CTRL |
| HNRNPR | 10236 | ENSP00000363741 | 12 | CTRL |
| RPS19 | 6223 | ENSP00000221975 | 12 | CTRL |
| VIM | 7431 | ENSP00000224237 | 12 | CTRL |
| TUBA1C | 84790 | ENSP00000301072 | 11 | CTRL |
| RPLP0 | 6175 | ENSP00000376299 | 11 | CTRL |
| ILF2 | 3608 | ENSP00000355011 | 11 | CTRL |
| RPL10 | 6134 | ENSP00000358832 | 11 | CTRL |
| HIST1H1E | 3008 | ENSP00000307705 | 11 | CTRL |
| PABPC1 | 26986 | ENSP00000313007 | 11 | CTRL |
| RPS15A | 6210 | ENSP00000318646 | 10 | CTRL |
| RPS7 | 6201 | ENSP00000339095 | 10 | CTRL |
| HNRNPA2B1 | 3181 | ENSP00000346694 | 10 | CTRL |
| HSP90AA1 | 3320 | ENSP00000216281 | 10 | CTRL |
| RPL4 | 6124 | ENSP00000311430 | 10 | CTRL |
| SLC25A5 | 292 | ENSP00000360671 | 10 | CTRL |
| ILF3 | 3609 | ENSP00000337305 | 10 | CTRL |
| RPS2 | 6187 | ENSP00000341885 | 9 | CTRL |
| RPL10A | 4736 | ENSP00000363018 | 9 | CTRL |
| RPS16 | 6217 | ENSP00000251453 | 9 | CTRL |
| RPS9 | 6203 | ENSP00000302896 | 9 | CTRL |
| HNRNPK | 3190 | ENSP00000365458 | 9 | CTRL |
| RPL15 | 6138 | ENSP00000309334 | 9 | CTRL |
| PARP1 | 142 | ENSP00000355759 | 9 | CTRL |
| HSPA1A | 3303 | ENSP00000364802 | 9 | CTRL |
| RPS8 | 6202 | ENSP00000379888 | 8 | CTRL |
| RPL22 | 6146 | ENSP00000346088 | 8 | CTRL |
| XRCC5 | 7520 | ENSP00000375978 | 8 | CTRL |
| XRCC6 | 2547 | ENSP00000353192 | 8 | CTRL |
| HNRNPL | 3191 | ENSP00000221419 | 8 | CTRL |
| TUBB4B | 10383 | ENSP00000341289 | 7 | CTRL |
| RPS24 | 6229 | ENSP00000361435 | 7 | CTRL |
| RPLP1 | 6176 | ENSP00000346037 | 7 | CTRL |
| NPM1 | 4869 | ENSP00000428755 | 7 | CTRL |
| RPLP2 | 6181 | ENSP00000322419 | 7 | CTRL |
| HNRNPD | 3184 | ENSP00000313199 | 7 | CTRL |
| HNRNPC | 3183 | ENSP00000338095 | 7 | CTRL |
| EEF1G | 1937 | ENSP00000331901 | 7 | CTRL |
| HIST1H1C | 3006 | ENSP00000339566 | 7 | CTRL |
| EEF1D | 1936 | ENSP00000434070 | 7 | CTRL |
| TUBB6 | 84617 | ENSP00000318697 | 7 | CTRL |
| RPL18A | 6142 | ENSP00000403914 | 6 | CTRL |
| PRPS1 | 5631 | ENSP00000361512 | 6 | CTRL |
| RPS13 | 6207 | ENSP00000228140 | 6 | CTRL |
| DARS | 1615 | ENSP00000264161 | 6 | CTRL |
| RPL14 | 9045 | ENSP00000345156 | 6 | CTRL |
| RPS18 | 6222 | ENSP00000393241 | 6 | CTRL |
| DDX5 | 1655 | ENSP00000440276 | 6 | CTRL |
| RPL21 | 6144 | ENSP00000346027 | 6 | CTRL |
| MRPS9 | 64965 | ENSP00000258455 | 6 | CTRL |
| TTN | 7273 | ENSP00000340554 | 6 | CTRL |
| RPS12 | 6206 | ENSP00000230050 | 5 | CTRL |
| RPL30 | 6156 | ENSP00000428085 | 5 | CTRL |
| MYL6 | 4637 | ENSP00000446955 | 5 | CTRL |
| RPS17 | 6218 | ENSP00000346045 | 5 | CTRL |
| GSTP1 | 2950 | ENSP00000381607 | 5 | CTRL |
| PRDX1 | 5052 | ENSP00000361152 | 5 | CTRL |
| RPS26 | 6231 | ENSP00000348849 | 5 | CTRL |
| THRAP3 | 9967 | ENSP00000346634 | 5 | CTRL |
| RPL8 | 6132 | ENSP00000378378 | 5 | CTRL |
| RPS10 | 6204 | ENSP00000347271 | 5 | CTRL |
| RPS6 | 6194 | ENSP00000369757 | 5 | CTRL |
| SNRPD1 | 6632 | ENSP00000300413 | 5 | CTRL |
| RPL27 | 6155 | ENSP00000253788 | 5 | CTRL |
| RPL11 | 6135 | ENSP00000363676 | 5 | CTRL |
| RPL12 | 6136 | ENSP00000354739 | 5 | CTRL |
| RPL18 | 6141 | ENSP00000447001 | 5 | CTRL |
| SNRNP70 | 6625 | ENSP00000221448 | 5 | CTRL |
| CFL1 | 1072 | ENSP00000432660 | 5 | CTRL |
| RPL27A | 6157 | ENSP00000346015 | 5 | CTRL |
| RPL13A | 23521 | ENSP00000375730 | 5 | CTRL |
| PDHB | 5162 | ENSP00000307241 | 5 | CTRL |
| HNRNPH1 | 3187 | ENSP00000377082 | 5 | CTRL |
| KARS | 3735 | ENSP00000325448 | 5 | CTRL |
| RPL13 | 6137 | ENSP00000307889 | 5 | CTRL |
| KRT1 | 3848 | K2C1_HUMAN | 27 | PieF_replicate_1 |
| HSP90AB1 | 3326 | ENSP00000360709 | 25 | PieF_replicate_1 |
| TUBB | 203068 | ENSP00000339001 | 21 | PieF_replicate_1 |
| TUBA1B | 10376 | ENSP00000336799 | 18 | PieF_replicate_1 |
| HSPD1 | 3329 | ENSP00000373620 | 18 | PieF_replicate_1 |
| KRT9 | 3857 | ENSP00000246662 | 17 | PieF_replicate_1 |
| HSPA8 | 3312 | ENSP00000437125 | 17 | PieF_replicate_1 |
| CLTC | 1213 | ENSP00000376763 | 16 | PieF_replicate_1 |
| KRT10 | 3858 | ENSP00000269576 | 15 | PieF_replicate_1 |
| EEF1A1 | 1915 | ENSP00000339063 | 14 | PieF_replicate_1 |
| MYH9 | 4627 | ENSP00000216181 | 12 | PieF_replicate_1 |
| KRT2 | 3849 | ENSP00000310861 | 12 | PieF_replicate_1 |
| NCL | 4691 | ENSP00000318195 | 11 | PieF_replicate_1 |
| RPL10 | 6134 | ENSP00000358832 | 11 | PieF_replicate_1 |
| ATP5B | 506 | ENSP00000262030 | 9 | PieF_replicate_1 |
| HSPA9 | 3313 | ENSP00000297185 | 9 | PieF_replicate_1 |
| HSPA5 | 3309 | ENSP00000324173 | 9 | PieF_replicate_1 |
| HSP90AA1 | 3320 | ENSP00000216281 | 9 | PieF_replicate_1 |
| HNRNPK | 3190 | ENSP00000365458 | 9 | PieF_replicate_1 |
| DHX9 | 1660 | ENSP00000356520 | 8 | PieF_replicate_1 |
| RPL10L | 140801 | ENSP00000298283 | 8 | PieF_replicate_1 |
| ATP5A1 | 498 | ENSP00000282050 | 7 | PieF_replicate_1 |
| FLNA | 2316 | ENSP00000353467 | 7 | PieF_replicate_1 |
| HNRNPA1 | 3178 | ENSP00000448617 | 7 | PieF_replicate_1 |
| CFL1 | 1072 | ENSP00000432660 | 7 | PieF_replicate_1 |
| RPL3 | 6122 | ENSP00000386101 | 6 | PieF_replicate_1 |
| RPL6 | 6128 | ENSP00000202773 | 6 | PieF_replicate_1 |
| NPM1 | 4869 | ENSP00000428755 | 6 | PieF_replicate_1 |
| GSTP1 | 2950 | ENSP00000381607 | 6 | PieF_replicate_1 |
| RPL7A | 6130 | ENSP00000361076 | 5 | PieF_replicate_1 |
| XRCC5 | 7520 | ENSP00000375978 | 5 | PieF_replicate_1 |
| PARP1 | 142 | ENSP00000355759 | 5 | PieF_replicate_1 |
| HNRNPC | 3183 | ENSP00000338095 | 5 | PieF_replicate_1 |
| KRT1 | 3848 | ENSP00000252244 | 14 | PieF_replicate_2 |
| TUBB | 203068 | ENSP00000339001 | 12 | PieF_replicate_2 |
| KRT10 | 3858 | ENSP00000269576 | 12 | PieF_replicate_2 |
| TUBA1B | 10376 | ENSP00000336799 | 9 | PieF_replicate_2 |
| RPL10 | 6134 | ENSP00000358832 | 9 | PieF_replicate_2 |
| KRT2 | 3849 | ENSP00000310861 | 8 | PieF_replicate_2 |
| ACTB | 60 | ENSP00000349960 | 6 | PieF_replicate_2 |
| HSPA8 | 3312 | ENSP00000437125 | 6 | PieF_replicate_2 |
| RPL10L | 140801 | ENSP00000298283 | 6 | PieF_replicate_2 |
| HSPD1 | 3329 | ENSP00000373620 | 5 | PieF_replicate_2 |
